# Characterizing the Metabolic Fate and Pharmacokinetics of the RyR2 Inhibitor *ent*-Verticilide

**DOI:** 10.64898/2026.10.31.763093

**Authors:** Tri Q. Do, Rachel D. Crouch, Madelaine P. Thorpe, Kevin M. Johnson, Jeanne Rumsey, Jeffrey N. Johnston, Björn C. Knollmann, Wendell S. Akers

## Abstract

*ent*-Verticilide is a synthetic macrocyclic depsipeptide and ryanodine receptor 2 (RyR2) inhibitor with potential therapeutic applications in cardiac arrhythmias. This study characterized its in vitro metabolism and in vivo pharmacokinetics to better understand factors that may influence its disposition. Metabolic stability was evaluated in mouse, rat, dog, and human liver S9 fractions, followed by CYP reaction phenotyping in human liver microsomes and metabolite profiling in human and rat liver S9 fractions. Pharmacokinetics were evaluated in rats following intravenous and oral administration. *ent*-Verticilide underwent rapid, NADPH-dependent depletion across all species, with in vitro half-lives ranging from 2.1 to 5.3 min. CYP reaction phenotyping indicated that CYP3A was the predominant CYP subfamily involved in its metabolism. Eight putative metabolites were identified in human and rat liver S9, primarily resulting from hydroxylation, oxidation, and desaturation, with generally similar metabolite profiles between species. In rats, *ent*-verticilide exhibited a systemic clearance of 29.6 mL/min/kg, a steady-state volume of distribution of 4.86 L/kg, and a terminal half-life of 6.85 h following intravenous administration. Absolute oral bioavailability was 10.5%. Overall, these findings show that *ent*-verticilide undergoes extensive oxidative hepatic metabolism and provide an initial DMPK framework for its continued preclinical development.

**SIGNIFICANCE STATEMENT:** This study provides the first integrated characterization of the metabolism and pharmacokinetics of *ent*-verticilide, a RyR2-targeting macrocyclic depsipeptide. *ent*-Verticilide was rapidly metabolized across species, with CYP3A identified as an important contributor to its oxidative metabolism. The compound also showed extensive distribution and low oral bioavailability in rats. These findings establish a foundation for understanding the disposition of *ent*-verticilide and highlight metabolic and pharmacokinetic factors that should be considered in its continued development.

## INTRODUCTION

*ent*-Verticilide is a synthetic 24-membered macrocyclic depsipeptide belonging to the cyclic oligomeric depsipeptide class.^1,2^ Unlike its naturally occurring enantiomer, *ent*-verticilide is available only through chemical synthesis and has been shown to potently inhibit ryanodine receptor 2 (RyR2).^1^ RyR2 regulates calcium release from the sarcoplasmic reticulum in cardiac myocytes,^3^ and dysregulated RyR2 activity can promote abnormal intracellular calcium handling and cardiac arrhythmias.^4^ Accordingly, RyR2 has emerged as a potential therapeutic target for cardiac arrhythmias.^4^

Previous studies from our laboratory demonstrated that, following intraperitoneal administration in mice, *ent*-verticilide exhibited a favorable pharmacokinetic profile and achieved systemic exposures sufficient to elicit pharmacodynamic activity and suppress arrhythmias.^5^ These findings supported *ent*-verticilide as a promising RyR2-targeting therapeutic candidate. However, further development requires a better understanding of its metabolic stability, biotransformation pathways, and pharmacokinetic properties across preclinical species. Such information is particularly important for a structurally complex macrocyclic compound, for which limited gastrointestinal absorption, hepatic metabolism, and extensive tissue distribution may influence systemic exposure.

The present study therefore aimed to characterize the in vitro metabolism and in vivo pharmacokinetics of *ent*-verticilide. Metabolic stability was evaluated in mouse, rat, dog, and human liver S9 fractions, and exploratory in vitro-to-in vivo scaling was used to estimate apparent hepatic intrinsic clearance and hepatic extraction. Reaction phenotyping with cytochrome P450-selective inhibitors was conducted in pooled human liver microsomes to identify the cytochrome P450 (P450) subfamilies contributing to parent-compound depletion. Metabolite profiling and tentative structural characterization were performed in human and rat liver S9 fractions using high-resolution tandem mass spectrometry to define the principal biotransformation pathways. Finally, systemic pharmacokinetics and absolute oral bioavailability were evaluated in rats following intravenous and oral administration.

## MATERIALS AND METHODS

### Chemicals and Reagents

Unless otherwise specified, chemicals and reagents were purchased from Sigma-Aldrich (St. Louis, MO, USA). *ent*-verticilide (batch#: MPT-4-274 with purity > 90%) was synthesized at Vanderbilt University (Nashville, TN, USA) using a previously reported synthetic procedure.^1^ Optima Liquid Chromatography/Mass Spectrometry (LC/MS)-grade acetonitrile and formic acid were obtained from Fisher Scientific (Waltham, MA, USA). Ultrapure water was generated using a Milli-Q Plus water-purification system (Millipore, Bedford, MA, USA). Potassium phosphate buffer components, magnesium chloride (MgCl₂), reduced nicotinamide adenine dinucleotide phosphate (NADPH), DMSO, carbamazepine, verapamil, montelukast, quinidine, and ketoconazole were obtained from Sigma-Aldrich (St. Louis, MO). α-Naphthoflavone and sulfaphenazole, were purchased from Santa Cruz Biotechnology (Dallas, TX). 2-phenyl-2-(1-piperidinyl)-propane (PPP) and (S)-(+)-N-3-benzylnirvanolwere purchased from Toronto Research Chemicals (Toronto, ON). Substrate (*ent*-verticilide, verapamil) stock solutions were prepared in DMSO, and working solutions were diluted into 1:1 water/acetonitrile (v/v). Cytochrome P450-selective inhibitor stock solutions were prepared in DMSO, and working solutions were diluted into potassium phosphate buffer (100 mM, pH 7.4). NADPH was prepared fresh in potassium phosphate buffer (100 mM, pH 7.4).

### Liver S9 Fractions and Human Liver Microsomes

For metabolic stability and reaction phenotyping studies, commercially available pooled liver S9 fractions from CD1 mouse (catalog number M1000.S9 / lot number 2110326 / n = 1400), Sprague-Dawley rat (catalog number R1000.S9 / lot number 2010058 / n = 97), Beagle dog (catalog number D1000.S9 / lot number 2110223 / n = 9), and human (catalog number H2620.S9 / lot number 1610440 / n = 200) were purchased from Sekisui XenoTech (Kansas City, KS). Human liver S9 fractions were pooled from a mixture of male and female donors. Liver S9 fractions from all other species were pooled from male donors. Pooled human liver microsomes (catalog number H2610 / lot number 1910096 / n = 200) were obtained from Sekisui XenoTech (Kansas City, KS, USA). Human liver microsomes were pooled from an equal mixture of male and female donors. The metabolite-profiling study was conducted by Inotiv, Inc. (Maryland Heights, MO, USA). Pooled, mixed-sex human liver S9 (catalog number H0610.S9 / lot number 1810001 / n = 50) was obtained from Sekisui XenoTech (Kansas City, MO, USA). Pooled male Sprague-Dawley rat liver S9 (catalog number M000012.S9 / lot number / n = 30) was obtained from BioIVT ADME-Tox (Baltimore, MD, USA). All liver S9 fractions and microsomes were stored at −80°C until use.

### Metabolic Stability of *ent*-Verticilide in Mouse, Rat, Dog, and Human Liver S9 Fractions

The metabolic stability of *ent*-verticilide was evaluated in pooled mouse, rat, dog, and human liver S9 fractions. Incubation mixtures were prepared in polypropylene microtubes with 100 mM potassium phosphate buffer, pH 7.4, containing 3 mM MgCl₂ and liver S9 at a final protein concentration of 2.5 mg/mL. The total incubation volume was 750 µL.

Incubation mixtures were preincubated with substrate for 15 minutes at 37°C. Metabolic reactions were initiated by adding NADPH (or vehicle for minus NADPH control samples) to a final concentration of 2 mM. Corresponding control incubations were conducted without NADPH. *ent*-Verticilide was evaluated at a final concentration of 1 µM. Verapamil was evaluated separately at 1 µM under otherwise identical conditions as a positive-control CYP3A substrate to verify NADPH-dependent metabolic activity. The final organic-solvent concentration was held constant within each experiment and did not exceed 0.6 %.

At 0, 3, 7, 15, 30, and 45 minutes, 100-µL aliquots were removed from each incubation and transferred immediately to a 96-well protein-precipitation filtration plate containing 300 µL of ice-cold acetonitrile supplemented with 50 nM carbamazepine as the internal standard. Precipitated proteins were removed by filtration into a polypropylene 96-well collection plate using a positive-pressure manifold. Filtered extracts were evaporated to dryness under a stream of nitrogen. The resulting residues were reconstituted in 100 µL of acetonitrile/water (30:70, v/v) before Liquid Chromatography Tandem Mass Spectrometry (LC–MS/MS) analysis.

### Reaction Phenotyping of *ent*-Verticilide in Human Liver Microsomes

The contributions of different P450 enzymes to the metabolism of *ent*-verticilide were evaluated in pooled human liver microsomes using P450-selective chemical inhibitors (**Supplemental Table S1**). *ent*-Verticilide (1 µM) was incubated at 37°C (shaking water bath) in a potassium phosphate buffered (100 mM, pH 7.4) mixture containing 3 mM MgCl_2_ and human hepatic microsomes (0.5 mg/ml), NADPH (2 mM) and the presence or absence of the P450-selective inhibitors. The final organic-solvent concentration was held constant within each experiment and did not exceed 0.6 %. Vehicle-control incubations contained an equivalent final concentration of DMSO without inhibitor. The total incubation volume was 300 µL.

Incubation mixtures were preincubated with substrate for 10 minutes at 37°C, followed by an additional 5-minute preincubation with the respective P450 inhibitor. Metabolic reactions were initiated by adding NADPH. At 0 and 60 minutes, 100-µL aliquots were removed from each incubation and transferred immediately to a 96-well protein-precipitation filtration plate containing 300 µL of ice-cold acetonitrile supplemented with 50 nM carbamazepine as the internal standard. Precipitated proteins were removed by filtration into a polypropylene 96-well collection plate using a positive-pressure manifold. Filtrates were evaporated to dryness under a stream of nitrogen. The resulting residues were reconstituted in 100 µL of acetonitrile/water (30:70, v/v) before LC–MS/MS analysis. Verapamil was evaluated separately as a positive-control CYP3A substrate at a final concentration of 1 µM under identical incubation conditions.

### Metabolite Profiling of *ent*-Verticilide in Rat and Human Liver S9 Fractions

The metabolite-profiling study was conducted by Inotiv, Inc. (Maryland Heights, MO, USA). Metabolite-profiling incubations were prepared in a 96-well plate fitted with 1-mL glass vial inserts. Separate incubation wells were prepared for the 0- and 60-minute time points. Each 100-µL incubation contained pooled human or rat liver S9 at a final protein concentration of 5.0 mg/mL in 100 mM potassium phosphate buffer, pH 7.4, containing 3 mM MgCl₂ and alamethicin at 50 µg/mg S9 protein. The final DMSO concentration was less than 0.25%. For each species, incubations were conducted either with both 2 mM NADPH and 2 mM uridine 5′-diphosphoglucuronic acid (UDPGA) or without either cofactor. Alamethicin was included in all incubations, including the no-cofactor controls. A liver S9 matrix blank without *ent*-verticilide was also prepared for each species. One incubation replicate was evaluated per species, cofactor condition, and time point.

Reactions were initiated by adding *ent*-verticilide to a final concentration of 25 µM. Samples were incubated at 37°C on a heated plate shaker operated at 500 rpm. At 0 or 60 minutes, the entire 100-µL sample from the corresponding incubation well was diluted with 100 µL of water and quenched with 600 µL of ice-cold acetonitrile supplemented with 50 ng/mL labetalol as the internal standard.

Samples were vortex-mixed and centrifuged at 1,000 × *g* for 10 minutes. The resulting supernatants were concentrated under a steady stream of nitrogen to a final volume of approximately 100 µL. Concentrated extracts were vortex-mixed, centrifuged briefly, and analyzed by High Performance Liquid Chromatography-High Resolution Tandem Mass Spectrometry (HPLC–HRMS/MS).

### In Vivo Pharmacokinetic Study in Rat

The pharmacokinetic study was conducted by the contract research organization Pharmaron, Inc. (Waltham, MA, USA). All study protocols and procedures were conducted in accordance with an Animal Care and Use Application approved by the Institutional Animal Care and Use Committee (IACUC) of Pharmaron, Inc. (protocol no. PH-DMPK-VUMC-22-001) and followed the guidance of the Association for Assessment and Accreditation of Laboratory Animal Care (AAALAC). Rats were housed under standard laboratory conditions with free access to food and water. Male Sprague–Dawley rats were 6–8 weeks of age and weighed approximately 223–252 g at the time of dosing. A total of 48 animals were included in the pharmacokinetic analysis, with 24 animals assigned to each route of administration.

The systemic pharmacokinetics of *ent*-verticilide were evaluated following single intravenous (IV) and oral (PO) administration to male Sprague–Dawley rats using a terminal destructive-sampling design. Plasma pharmacokinetic sampling was incorporated as a substudy within a broader exploratory *in vivo* study requiring terminal sample collection. The present analysis was restricted to systemic plasma concentrations of *ent*-verticilide. *ent*-Verticilide was administered as a single 3 mg/kg IV dose via the lateral tail vein or as a single 9 mg/kg PO dose by oral gavage. For both routes, the dosing vehicle consisted of 10% Tween 20, 10% dimethyl sulfoxide (DMSO), 40% water, and 40% polyethylene glycol 400 (PEG400). Final dosing concentrations were 0.6 mg/mL for IV administration and 0.9 mg/mL for PO administration, corresponding to nominal dose volumes of 5 and 10 mL/kg, respectively. Individual dosing volumes were adjusted according to animal body weight, and dosing formulations were maintained at room temperature until administration. A terminal destructive-sampling design was used, with three animals assigned to each of eight pharmacokinetic sampling times for each route of administration. Blood samples were collected at 0.083, 0.167, 0.5, 1, 3, 5, 10, and 24 h post dose. At each scheduled time point, approximately 200 µL of blood was collected by cardiac puncture into tubes containing K₂EDTA as the anticoagulant. Plasma was separated by centrifugation at 4000 × *g* for 5 min at 4°C and stored at −80 ± 15°C until bioanalysis. Because separate animals were sampled at each time point, each animal contributed a single terminal pharmacokinetic blood sample rather than undergoing serial blood sampling.

### LC–MS/MS Analysis

#### Metabolic Stability and Reaction Phenotyping Experiments

Samples were introduced using a CTC Analytics PAL-xt DW autosampler, and 10 µL was injected into a Shimadzu LC-20AD series HPLC system coupled to a SCIEX 6500 QTRAP mass spectrometer (SCIEX, Foster City, CA, USA). The mass spectrometer was equipped with an electrospray ionization source (500°C ion source temperature) operated in positive-ion mode (4500 V ion spray voltage). Chromatographic separation was performed using a Kinetex® C18 column (50 × 2.1 mm, 1.7-µm particle size; Phenomenex, Torrance, CA, USA) maintained at 60°C. Mobile phase A consisted of 0.1% formic acid in ultrapure water, and mobile phase B consisted of 0.1% formic acid in acetonitrile. The mobile phases were delivered at a flow rate of 0.5 mL/min. The initial mobile-phase composition was 5% B. Mobile phase B was linearly increased from 5% to 90% over 0.5 minutes. The 90% B condition was maintained from 0.5 to 2.00 minutes, after which the composition was returned to 5% B from 2.00 to 2.01 minutes. The column was re-equilibrated at 5% B until the end of the 2.5-minute analytical run.

*ent*-Verticilide was detected by multiple-reaction monitoring using the precursor-to-product-ion transition *m/z* 853.6→214.1. Carbamazepine (internal standard) was monitored using the transition *m/z* 237.1→194.0. Verapamil was monitored using the transition *m/z* 455.0→165.0. Peak areas were integrated using Analyst 1.7.0. The ion-source parameters, collision energies, and MRM dwell times for *ent*-verticilide, carbamazepine, and verapamil are reported in **Supplemental Table S2**.

#### Metabolite Profiling of *ent*-Verticilide in Rat and Human Liver S9 Fractions

Metabolite profiling and tentative structural characterization were performed using HPLC–HRMS/MS. Mass-spectrometric analysis was performed using an Orbitrap Exploris 120 mass spectrometer (Thermo Fisher Scientific, Waltham, MA, USA) equipped with a heated electrospray ionization source operated in positive-ion mode. The spray voltage was 3.5 kV, and the RF lens setting was 70%. The ion-transfer-tube and vaporizer temperatures were 325°C and 350°C, respectively. Sheath, auxiliary, and sweep gas settings were 50, 10, and 1 arbitrary units, respectively. Chromatographic separation was performed using a Vanquish Flex HPLC system equipped with a Vanquish Flex autosampler (Thermo Fisher Scientific, Waltham, MA, USA) and an Restek Ultra C8 column (2.1 × 150 mm, 3-µm particle size). The column was maintained at 40°C, and the autosampler was maintained at 4°C. Mobile phase A consisted of 0.1% formic acid in ultrapure water, and mobile phase B consisted of 0.1% formic acid in acetonitrile. The mobile phases were delivered at 0.4 mL/min. The gradient was maintained at 5% B from initialization through 2 minutes, increased linearly to 80% B from 2 to 8 minutes, and then increased to 95% B from 8 to 14 minutes. The composition was maintained at 95% B from 14 to 16.5 minutes, returned to 5% B from 16.5 to 17 minutes, and maintained at 5% B through 20 minutes for column re-equilibration. The injection volume was 10 µL, and the total chromatographic run time was 20 minutes.

Full-scan high-resolution mass spectra were acquired over an *m/z* range of 150–1500 at a resolution of 60,000. Data-dependent MS/MS spectra were acquired at a resolution of 15,000 using stepped normalized collision-energy settings of 15, 25, and 35. The reported product-ion spectra combined MS/MS data acquired across all three stepped collision-energy settings. Data acquisition was performed using Xcalibur version 4.4.16.14. Chromatographic and mass-spectral data were reviewed using FreeStyle version 1.8.63.0 (Thermo Fisher Scientific, Waltham, MA, USA).

#### In Vivo Pharmacokinetic Study

Plasma concentrations of *ent*-verticilide were quantified using a qualified HPLC–MS/MS assay. Chromatographic separation was performed using a Shimadzu LC-30AD Series HPLC system equipped with a SIL-30AC autosampler (Shimadzu Corporation, Kyoto, Japan). Analytes were separated on a Raptor Biphenyl column (50 × 2.1 mm, 2.7-µm particle size; Restek Corporation, Bellefonte, PA, USA). Mobile phase A consisted of 0.1% formic acid in ultrapure water, and mobile phase B consisted of 0.1% formic acid in acetonitrile–water (95:5, v/v). The mobile phases were delivered at a flow rate of 0.6 mL/min. The initial composition of 10% B was maintained for 0.2 min. Mobile phase B was then increased linearly from 10% to 100% between 0.2 and 1.4 min and maintained at 100% until 1.90 min. The mobile-phase composition was returned to 10% B between 1.91 and 2.30 min and maintained at 10% B until 2.50 min to re-equilibrate the column before the subsequent injection. Analytes were detected using an AB SCIEX API 4000 mass spectrometer equipped with an electrospray-ionization source operated in positive-ion mode. *ent*-Verticilide and dexamethasone, the internal standard, were monitored using the precursor-to-product ion transitions *m/z* 853.6→214.1 and 393.4→373.3, respectively.

Calibration curves were constructed by plotting the *ent*-verticilide-to-internal-standard peak-area ratio against the nominal *ent*-verticilide concentration. Study-sample concentrations were calculated using weighted linear regression with a weighting factor of 1/*x*². An analytical run was accepted when the calibration curve contained at least five standards with back-calculated concentrations within ±15% of their nominal values and a coefficient of determination (*r*) greater than 0.95. At least 66.7% of all quality-control (QC) samples and at least 50% of the QC samples at each concentration were required to be within ±15% of their nominal values.

### Data Analysis

#### Metabolic-Stability Analysis

For each sample, the LC–MS/MS area ratio was calculated by dividing the analyte peak area by the carbamazepine peak area:

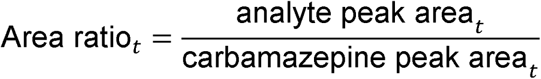

The percentage of *ent*-verticilide or verapamil remaining at each time point was calculated by normalizing the area ratio at that time point to the corresponding area ratio at 0 minutes:

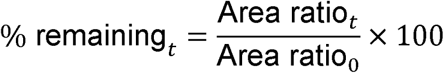

The percentage of parent compound remaining was natural-log transformed and plotted against incubation time. The first-order depletion rate constant, *k*, was determined as the negative of the slope of the log-linear regression:

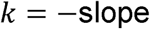

The in vitro half-life was calculated as:

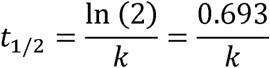

Measurements below 5% remaining substrate were excluded from regression analysis.

*In vitro* hepatic intrinsic clearance (*CL_int_*_,*app*,*scaled*_, ml/min/kg) for each species was estimated from hepatic S9 using the substrate depletion method according to the equation below:^6^

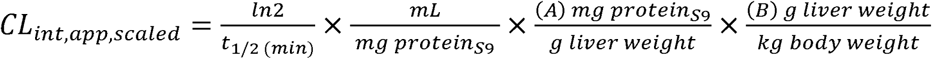

An S9 value (mg protein per g liver weight) of A = 120.7 was used for calculations in all species, as previously reported.^6,7^ Values for B (g liver weight per kg body weight) were also derived from the literature; B = 25, 87.5, 40, and 32 for human, mouse, rat, and dog, respectively.^8^

Because binding within the liver S9 incubations was not incorporated into the calculations, the resulting values were designated scaled apparent intrinsic clearance. Binding-unadjusted hepatic clearance was predicted using the well-stirred model:^9^

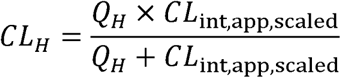

The corresponding hepatic extraction ratio was calculated as:

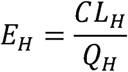

Species-specific hepatic blood-flow values of 90, 70, 31, and 21 mL/min/kg were used for mouse, rat, dog, and human, respectively.^8,10^

These calculations did not incorporate binding in plasma, blood, or the liver S9 incubations. Accordingly, they were considered exploratory, binding-unadjusted predictions rather than definitive estimates of in vivo hepatic clearance. In the absence of binding terms, the model is mathematically equivalent to assuming that the entire measured drug concentration was available for hepatic metabolism.

Although a blood-to-plasma concentration ratio of 0.85 was reported previously for *ent*-verticilide,^5^ that value was not incorporated because blood and plasma binding corrections were outside the scope of the current exploratory analysis.

### Reaction Phenotyping in Human Liver Microsomes

For the reaction-phenotyping experiments, the percentage of *ent*-verticilide remaining after 60 minutes was calculated by normalizing the analyte-to-internal-standard peak-area ratio at 60 minutes to the corresponding ratio at 0 minutes:

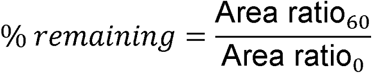

### Metabolite Profiling of *ent*-Verticilide in Rat and Human Liver S9 Fractions

An *ent*-verticilide reference standard was used to establish the accurate mass, chromatographic retention time, and characteristic MS/MS fragmentation pattern of the parent compound. These data served as references for metabolite detection and tentative structural characterization. Full-scan HRMS data were interrogated for potential metabolites of *ent*-verticilide. Candidate metabolites were identified by comparing chromatographic and mass-spectral signals in the 60-minute incubation samples with those in the corresponding 0-minute samples, species-specific matrix blanks, and no-cofactor controls. Proposed biotransformations were evaluated using accurate mass, predicted elemental composition, mass shift relative to *ent*-verticilide, chromatographic retention time, characteristic product ions, comparison of metabolite MS/MS fragmentation patterns with those of the parent compound, and predicted metabolic pathways. Metabolite-data analysis and structural evaluation were performed using Xcalibur, FreeStyle, Mass-MetaSite (Mass Analytica), and WebMetabase (Mass Analytica). Structural assignments for the detected metabolites were based on exact mass measurements and characterization of product-ion MS/MS fragments.

### Accurate-Mass Analysis

Mass error was calculated in parts per million using the observed and theoretical protonated molecular masses:

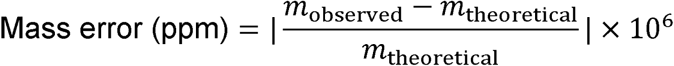

### Relative-Abundance Analysis

The relative abundance of *ent*-verticilide, putative metabolites M1–M8, and the C6 isomeric contaminant was calculated using LC–MS peak areas:

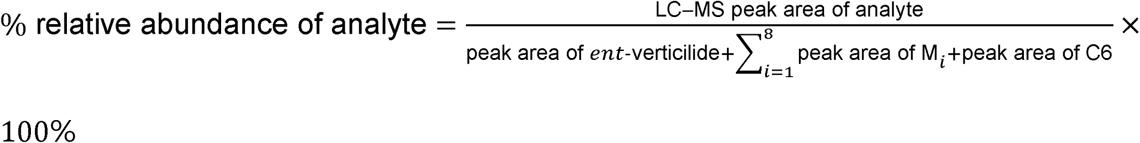

Chromatographic impurities C1–C5 were excluded from the denominator. C6 was included because it represented a chromatographically resolved isomeric contaminant with the same accurate mass as *ent*-verticilide and was considered part of the profiled test-article composition. Analytes representing less than 0.1% of the total included LC–MS peak area were reported as not detected (ND).

Relative LC–MS peak areas were used as a semiquantitative metric for metabolite-profile composition. Relative abundance values were not corrected for extraction recovery, ionization efficiency, adduct formation, and detector response due to the absence of authentic standards. Consequently, these relative abundance values represent single-incubation results and function strictly as qualitative, exploratory comparisons between human and rat S9 fractions rather than absolute concentrations.

### Pharmacokinetic Study

Plasma *ent*-verticilide concentrations at each nominal sampling time were summarized as the arithmetic mean, standard deviation (SD), and coefficient of variation (CV%) for three animals. Because a terminal destructive-sampling design was used, different animals contributed samples at each time point and individual longitudinal concentration–time profiles were not available.

Pharmacokinetic parameters were estimated using the sparse-sampling noncompartmental analysis (NCA) functionality in Phoenix WinNonlin version 8.5.2.4 (Certara, Princeton, NJ, USA). Individual plasma concentration measurements from each animal were entered at the corresponding nominal sampling time, with animal identification retained as the subject identifier. Separate NCA analyses were performed for the IV and PO treatment groups using the plasma concentration models for intravenous bolus and extravascular administration, respectively.

For sparse-sampling analysis, Phoenix calculated the arithmetic mean plasma concentration at each nominal sampling time and constructed a composite mean concentration–time profile for each route of administration. Pharmacokinetic parameters were derived from these composite profiles. Following PO administration, the maximum observed plasma concentration (*C_max_*) and time to maximum concentration (*T_max_*) were obtained directly from the composite concentration–time profile. For IV administration, the concentration at time zero (*C*_0_) was estimated by back-extrapolation according to the IV bolus NCA model. The terminal elimination rate constant (*λ_z_*) was estimated by log-linear regression of the terminal portion of the composite concentration–time profile, and terminal elimination half-life (*T*_1/2_) was calculated as 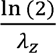. AUC extrapolated to infinity (*AUC_inf_*) was calculated using the observed last quantifiable concentration and *λ_z_*. The percentage of *AUC_inf_* attributable to extrapolation beyond the final sampling time (*AUC* extrapolated) was also determined.

Following IV administration, systemic clearance (*CL*) was calculated from dose and *AUC_inf_*. Steady-state volume of distribution (*V_ss_*) was determined using standard noncompartmental methods. Following PO administration, apparent oral clearance and apparent terminal volume of distribution were calculated by Phoenix but were not emphasized because IV administration permitted direct estimation of systemic clearance, distribution, and absolute oral bioavailability.

Absolute oral bioavailability (*F*) was calculated from dose-normalized *AUC_inf_* according to:

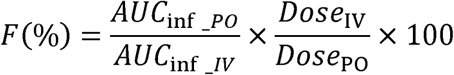

using IV and PO doses of 3 and 9 mg/kg, respectively.

Because pharmacokinetic parameters were derived from composite mean concentration–time profiles rather than serial measurements obtained from individual animals, the resulting NCA parameters represent group-level pharmacokinetic estimates. Accordingly, derived pharmacokinetic parameters were reported as point estimates rather than as mean ± SD. Interanimal variability in observed plasma concentrations at each nominal sampling time was summarized using SD and CV%.

## RESULTS

### NADPH-Dependent Metabolic Stability Across Species

*ent*-Verticilide underwent rapid, time-dependent depletion in mouse, rat, dog, and human liver S9 fractions in the presence of NADPH (**Supplemental Figure S1**). In contrast, little or no depletion was observed in the corresponding incubations without NADPH, supporting a predominant contribution of NADPH-dependent enzymatic metabolism to parent-compound depletion under the evaluated conditions. Verapamil also underwent time-dependent depletion in the presence of NADPH and remained comparatively stable in its absence, confirming the metabolic competence of the liver S9 preparations (data not shown).

The estimated first-order depletion-rate constants, in vitro half-lives, scaled apparent hepatic intrinsic-clearance values, and exploratory predictions of hepatic clearance and extraction ratio are summarized in **Table 1**. These exploratory estimates indicate high predicted hepatic extraction in all four species under the assumptions of the binding-unadjusted model.

**Table 1:**
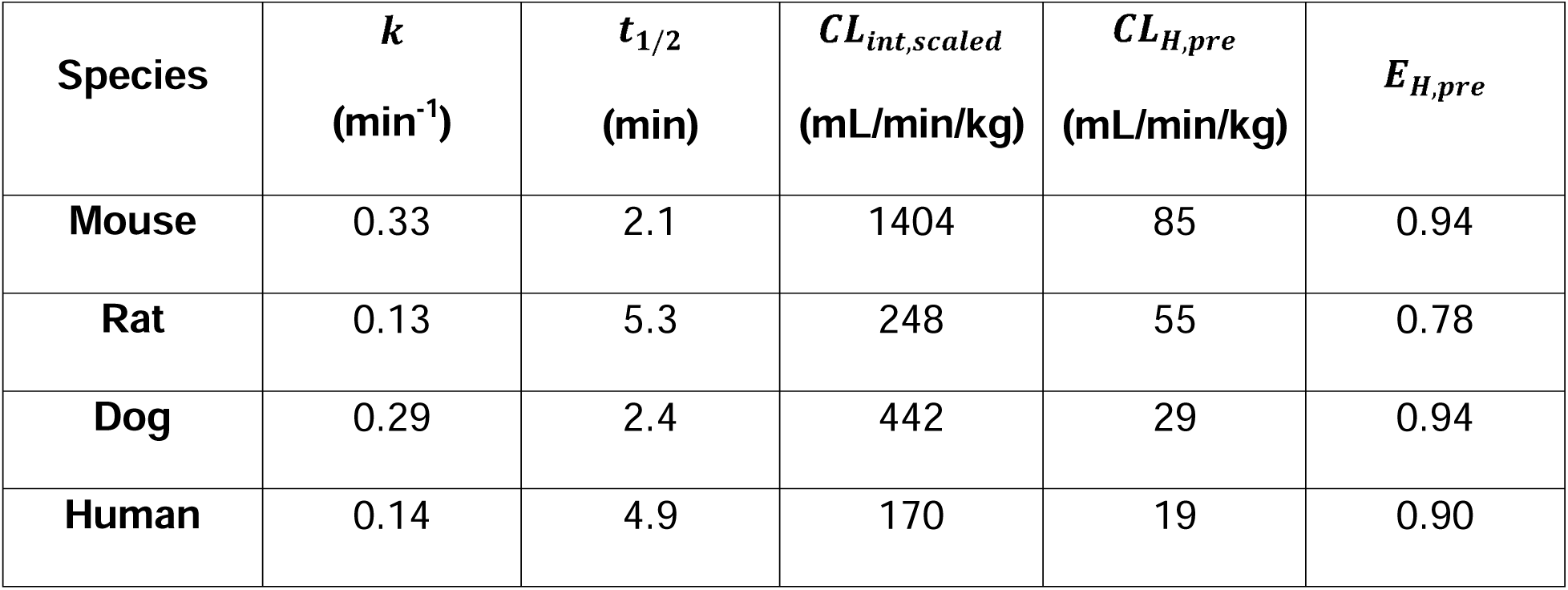
Predicted Parameters from Substrate Depletion of *ent*-Verticilide in Mouse, Rat, Dog, and Human Liver S9. The parameters were calculated using the equations described in the method section. ***k***: First-order elimination rate constant; ***t*_1_**_/**2**_: In vitro half-life; ***CL_int_***_,***scaled***_: Scaled hepatic intrinsic clearance; ***CL_H_***_,***pre***_: Predicted hepatic clearance; ***E_H_***_,***pre***_: Predicted hepatic extraction ratio.

### Reaction Phenotyping in Human Liver Microsomes

P450-selective inhibitors produced markedly different effects on the percentage of *ent*-verticilide remaining after incubation in pooled human liver microsomes (**Figure 1**). Ketoconazole, a CYP3A4/5 inhibitor, produced the greatest increase in the percentage of *ent*-verticilide remaining compared with the vehicle-control incubation.

**Figure 1:**
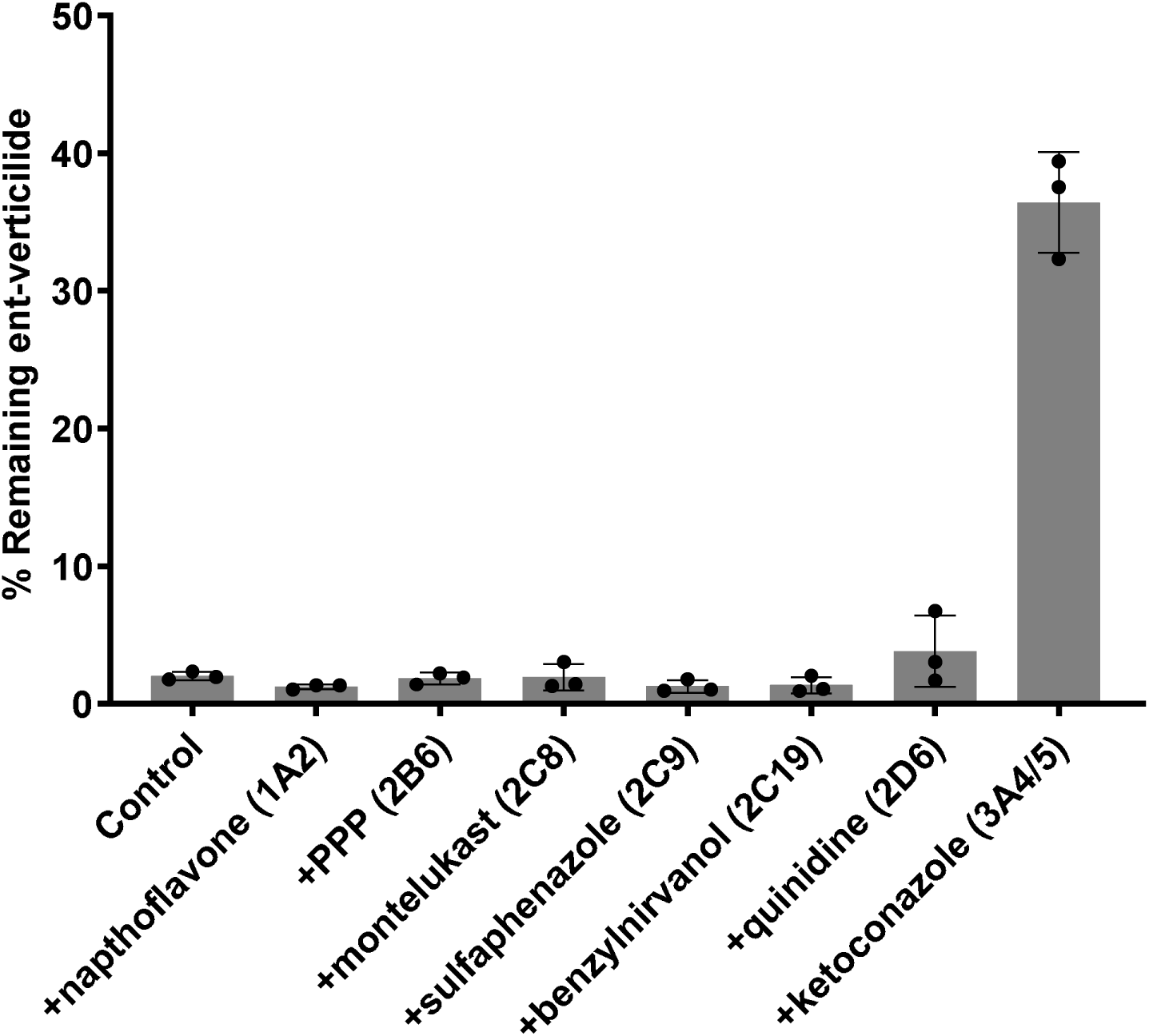
*ent*-Verticilide is Predominantly Metabolized by CYP3A4/5. X-axis represents the specific inhibitor and its corresponding target P450 isoform in parenthesis. Y-axis represents the percentage *ent*-verticilide (substrate) remaining after 60-minute incubation. *ent*-Verticilide (1 mM) was incubated at 37°C (shaking water bath) in a potassium phosphate buffered (100 mM, pH 7.4) mixture containing 3 mM MgCl_2_ and human hepatic microsomes (0.5 mg/ml), NADPH (2 mM) and the presence or absence of the P450-selective inhibitors. Verapamil, a CYP3A4 substrate, was included as a positive control. Data = mean ± S.D. from a single experiment performed in triplicate. **PPP**: 2-phenyl-2-(1-piperidinyl)propane.

Ketoconazole also increased the percentage of verapamil remaining, supporting effective inhibition of CYP3A activity under the assay conditions (data not shown). Inhibitors nominally selective for CYP1A2, CYP2B6, CYP2C8, CYP2C9, CYP2C19, and CYP2D6 did not increase the percentage of *ent*-verticilide remaining to an extent comparable with ketoconazole.

### Identification of Putative *ent*-Verticilide Metabolites in Human and Rat Liver S9 Fractions

HPLC–HRMS/MS analysis identified eight putative metabolites of *ent*-verticilide, designated M1–M8 (**Figure 2**). Protonated *ent*-verticilide was observed at *m/z* 853.5533 with a retention time of approximately 12.3 minutes. An isomeric contaminant designated C6 had the same accurate mass and eluted at approximately 12.6 minutes.

**Figure 2.**
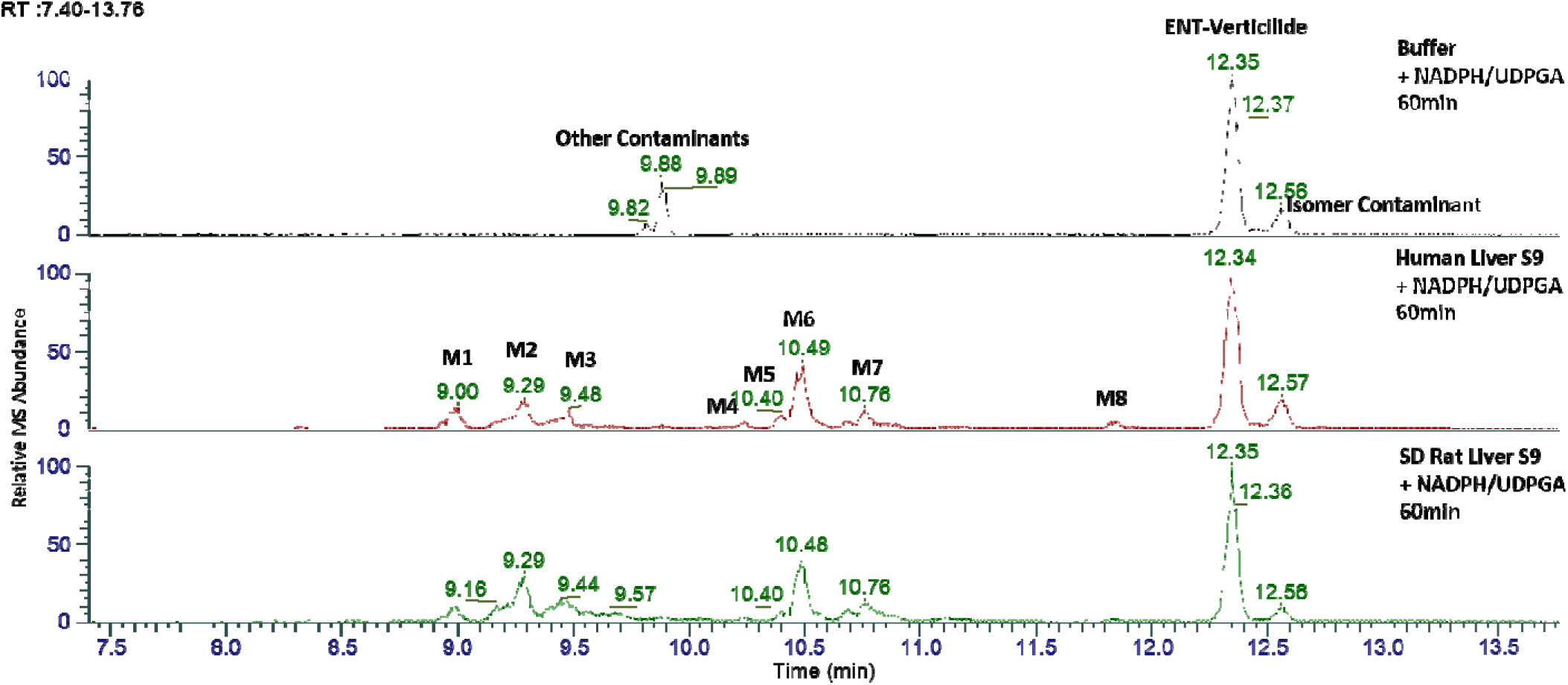
LCMS Chromatograms of *ent*-Verticilide and Metabolites in Liver S9 Fractions. Representative LC–MS chromatograms of *ent*-verticilide incubated with buffer (top), human liver S9 fractions (middle), or Sprague–Dawley (SD) rat liver S9 fractions (bottom) in the presence of NADPH and UDPGA for 60 min. Chromatographic peak intensities are expressed as relative MS abundance.

Prominent ammonium (M+18) and sodium (M+23) adducts were observed for the parent macrocycle and its putative metabolites. M1 and M2 exhibited mass increases of approximately 32 Da relative to *ent*-verticilide, consistent with dihydroxylation. M3 and M4 exhibited mass increases of approximately 30 Da, consistent with combined hydroxylation and oxidation. M5 and M6 exhibited mass increases of approximately 16 Da, consistent with monohydroxylation. Although M5 and M6 had similar retention times, the peaks were sufficiently resolved chromatographically to permit separate peak-area integration. M7 exhibited a mass increase of approximately 14 Da, consistent with oxidation, whereas M8 exhibited a mass decrease of approximately 2 Da, consistent with desaturation. Accurate-mass data are summarized in **Supplemental Table S3**, and full-scan MS and MS/MS spectra for *ent*-verticilide and M1–M8 are presented in **Supplemental Figures S2–S10**.

Relative-abundance values for S9 incubations were calculated using a denominator comprising *ent*-verticilide, isomer C6, and M1–M8 (**Table 2**). In human liver S9 incubations containing NADPH and UDPGA, the relative abundance of *ent*-verticilide decreased from 76.1% at 0 minutes to 42.6% at 60 minutes, whereas the relative abundance of C6 decreased from 23.9% to 8.6%. All eight putative metabolites were detected at 60 minutes.

**Table 2:** Relative LCMS Peak Areas of *ent*-Verticilide and Putative Metabolites.

| **Analyte** | **Proposed Biotransformation** | **Human 0 min** | **Human 60 min** | **Rat 0 min** | **Rat 60 min** |
|---|---|---|---|---|---|
| M1 | Dihydroxylation | ND | 5.3% | ND | 3.9% |
| M2 | Dihydroxylation | ND | 9.6% | ND | 17.4% |
| M3 | Hydroxylation and oxidation | ND | 5.2% | ND | 9.4% |
| M4 | Hydroxylation and oxidation | ND | 1.4% | ND | 0.9% |
| M5 | Hydroxylation | ND | 2.7% | ND | 1.9% |
| M6 | Hydroxylation | ND | 17.7% | ND | 18.5% |
| M7 | Oxidation | 0.1% | 4.9% | 0.1% | 6.6% |
| M8 | Desaturation | ND | 1.8% | ND | 0.6% |
| *ent*-Verticilide | Parent compound | 76.1% | 42.6% | 74.3% | 37.4% |
| C6 | Isomeric contaminant | 23.9% | 8.6% | 25.7% | 3.6% |

Monohydroxylation, M6, was the most abundant metabolite signal at 17.7%, followed by M2 (9.6%), M1 (5.3%), M3 (5.2%), M7 (4.9%), M5 (2.7%), M8 (1.8%), and M4 (1.4%). In rat liver S9 incubations containing NADPH and UDPGA, the relative abundance of *ent*-verticilide decreased from 74.3% at 0 minutes to 37.4% at 60 minutes, whereas the relative abundance of C6 decreased from 25.7% to 3.6%. All eight putative metabolites were detected at 60 minutes. M6 and M2 were the most abundant metabolite signals at 18.5% and 17.4%, respectively, followed by M3 (9.4%), M7 (6.6%), M1 (3.9%), M5 (1.9%), M4 (0.9%), and M8 (0.6%). In the buffer-control incubation containing NADPH and UDPGA in the absence of liver S9, M1–M5 and M7–M8 were below the 0.1% reporting threshold at both 0 and 60 minutes. M6 accounted for 0.1% of the total LC–MS peak area at both time points (Data not shown). The relative abundances of *ent*-verticilide and C6 were 82.1% and 17.8%, respectively, at 0 minutes and 87.6% and 12.4%, respectively, at 60 minutes. The absence of substantive metabolite formation in this control supports the requirement of liver S9 matrix for formation of M1–M8 under the evaluated conditions.

Based on the observed mass shifts, MS/MS fragmentation patterns, and predicted biotransformations, a tentative metabolic pathway was proposed for *ent*-verticilide in human and rat liver S9 fractions (**Figure 3**). The predominant in vitro metabolic pathway in both human and rat liver fractions was proposed to be hydroxylation and oxidation of the alkyl side chains. No evidence of hydrolysis or N-demethylation was observed and none of the metabolites M1-

**Figure 3.**
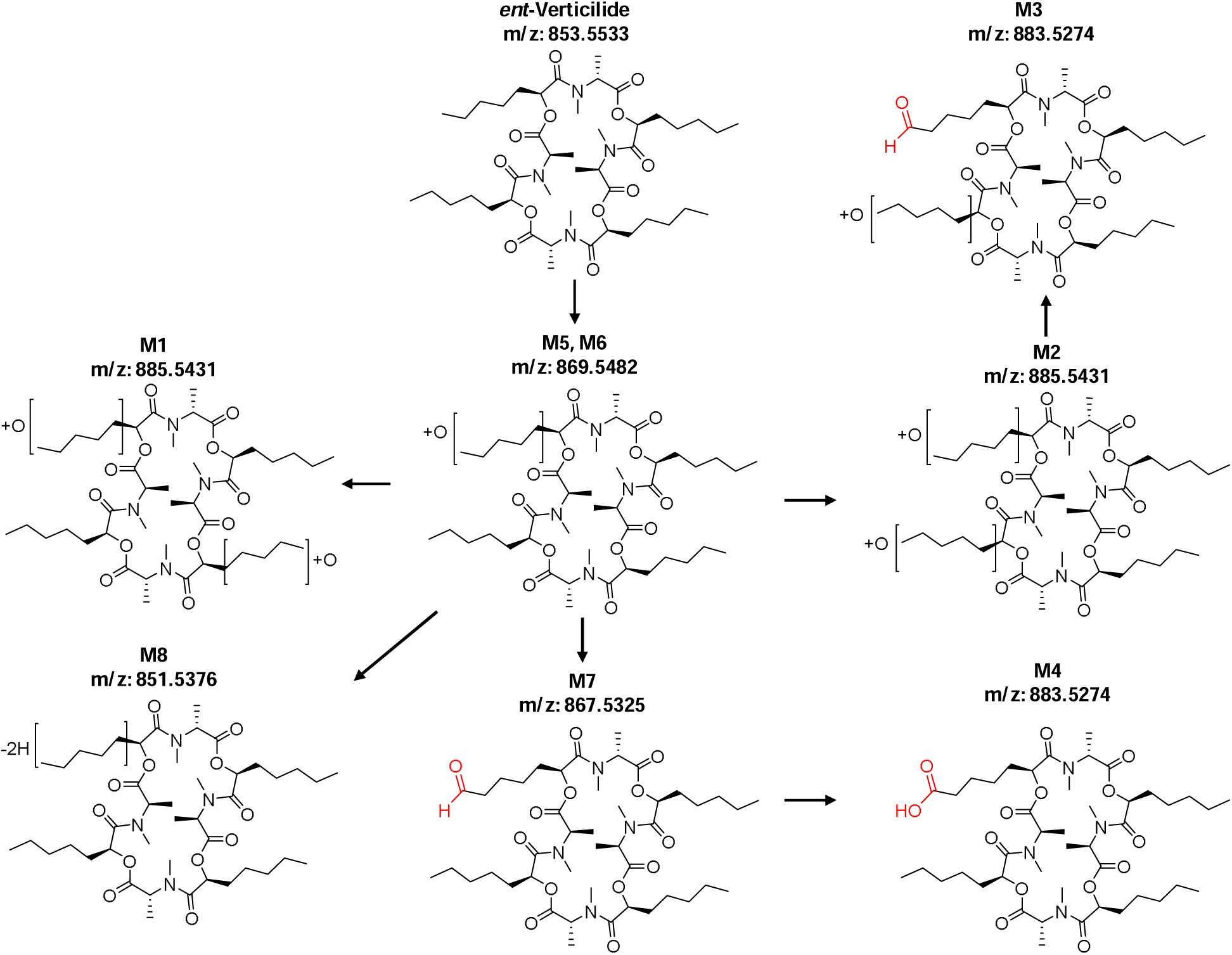
Proposed Metabolic Pathway of ent-Verticilide in Human and Rat Liver S9.

M8 exhibited a signal consistent with a glucuronide conjugate, despite the inclusion of UDPGA. Furthermore, no unique metabolites were identified in human liver fractions. In the absence of authentic metabolite standards, the proposed biotransformations and structures were considered tentative.

### In Vivo Pharmacokinetics of *ent*-Verticilide in Rats

Following a single 3-mg/kg intravenous dose, *ent*-verticilide exhibited a rapid initial decline in plasma concentration followed by a slower terminal phase (**Figure 4**). Systemic clearance and steady-state volume of distribution were

**Figure 4.**
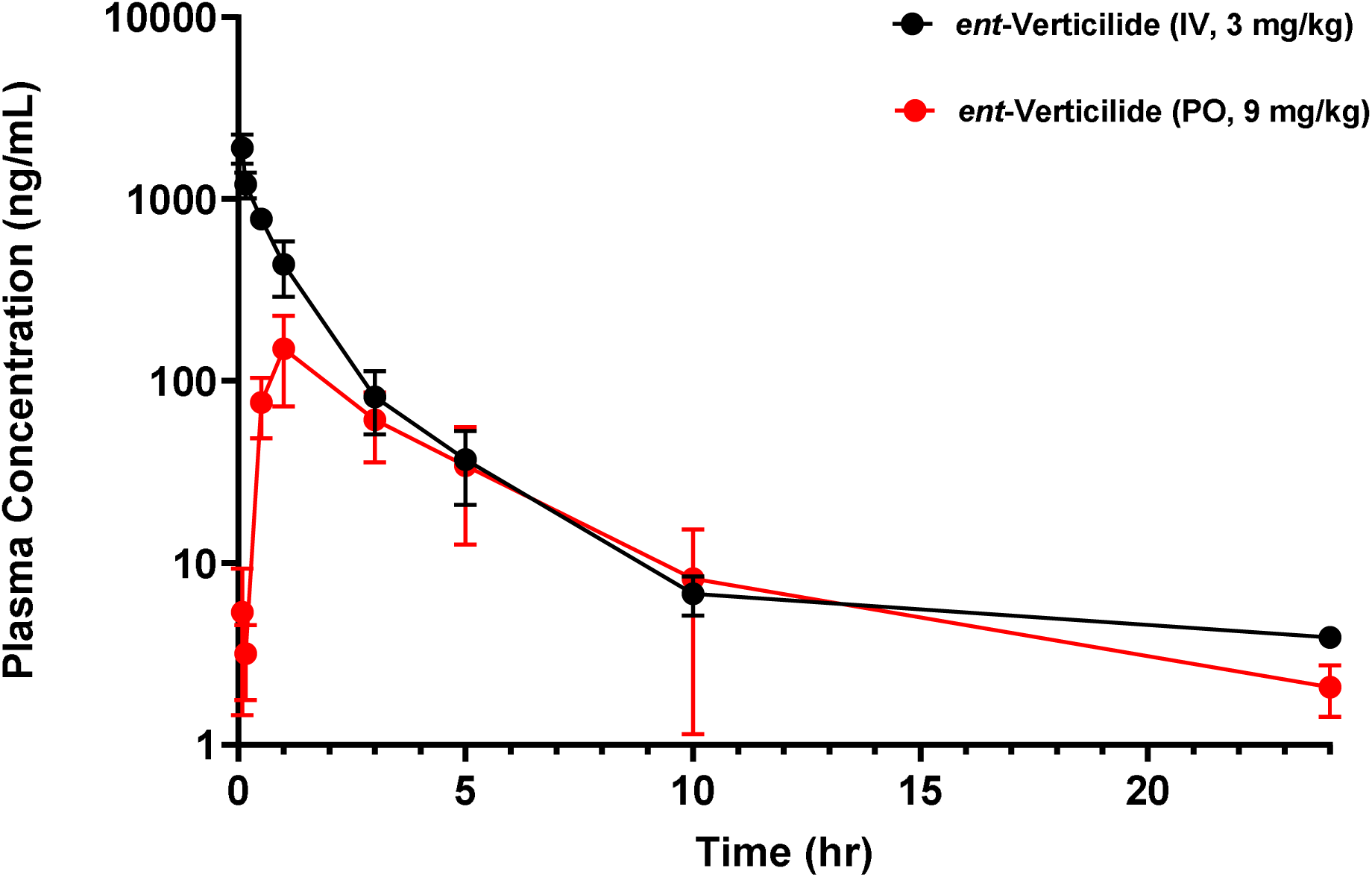
Plasma Concentration–Time Profiles of *ent*-Verticilide Following Intravenous and Oral Administration to Male Sprague–Dawley Rats. Mean plasma concentrations of *ent*-verticilide following a single 3 mg/kg intravenous (IV) dose or 9 mg/kg oral (PO) dose are shown through 24 h postdose. Data are presented as mean ± SD (n = 3 animals per nominal sampling time) using a terminal destructive-sampling design. Separate animals contributed plasma samples at each sampling time. Plasma concentrations are displayed using a logarithmic y-axis.

29.6 mL/min/kg and 4.86 L/kg, respectively, and the terminal elimination half-life was 6.85 hours (**Table 3**).

**Table 3.** Plasma Pharmacokinetic Parameters of *ent*-Verticilide Following Single Intravenous (IV) and Oral (PO) Administration to Male Sprague–Dawley Rats. Pharmacokinetic parameters represent group-level estimates derived from composite mean plasma concentration–time profiles using sparse-sampling noncompartmental analysis. Three animals contributed independent plasma samples at each nominal sampling time. *C*_0_ represents the back-extrapolated plasma concentration immediately following IV administration. Absolute oral bioavailability (***F***) was calculated from dose-normalized *AUC_inf_*. Derived pharmacokinetic parameters are point estimates and do not represent mean values from individual animals. ***C_max_***: The maximum observed plasma concentration; ***T_max_***: Time to maximum concentration; ***C*_0_**: The concentration at time zero; ***T*_1_**_/**2**_: Terminal elimination half-life; ***AUC_inf_***: The area under the concentration-versus-time curve extrapolated to infinity; ***CL***: Systemic clearance; ***V_ss_***: Steady-state volume of distribution.

| **Pharmacokinetic Parameter** | **IV, 3 mg/kg** | **PO, 9 mg/kg** |
|---|---|---|
| *C*<sub>0</sub> (ng/mL) | 3015 | — |
| *C*<sub>*max*</sub> (ng/mL) | — | 150 |
| *T*<sub>*max*</sub> (h) | — | 1.0 |
| *AUC*<sub>*inf*</sub> (ng·h/mL) | 1687 | 530 |
| *AUC* extrapolated (%) | 2.28 | 2.87 |
| *T*<sub>1/2</sub> (h) | 6.85 | 5.07 |
| *CL* (mL/min/kg) | 29.6 | — |
| *V*<sub>*ss*</sub> (L/kg) | 4.86 | — |
| *F* (%) | — | 10.5 |

Intravenous exposure was characterized by an estimated initial plasma concentration of 3015 ng/mL and an AUC extrapolated to infinity of 1687 ng·h/mL. Following a single 9-mg/kg oral dose, plasma concentrations increased during the first hour after administration and reached an observed maximum concentration of 150 ng/mL at an observed time to maximum concentration of 1 hour. Concentrations subsequently declined through 24 hours (**Figure 4**). The estimated terminal elimination half-life following oral administration was 5.07 hours. Following dose normalization for both routes of administration, the estimated absolute oral bioavailability was approximately 10.5%.

## DISCUSSION

The present studies characterized the metabolic stability, P450 reaction phenotype, metabolite profile, and rat pharmacokinetics of *ent*-verticilide. Across mouse, rat, dog, and human liver S9 fractions, *ent*-verticilide underwent rapid, predominantly NADPH-dependent depletion. Reaction-phenotyping experiments in pooled human liver microsomes implicated CYP3A4/5 as the principal P450 enzymes contributing to this metabolism. Consistent with these observations, metabolite-profiling experiments identified several putative oxidative metabolites in human and rat liver S9 fractions. Together, these findings indicate that oxidative hepatic metabolism is an important component of the disposition of *ent*-verticilide and may contribute to its limited oral exposure.

Exploratory application of the binding-unadjusted well-stirred model predicted high hepatic extraction in all four species. These estimates should be interpreted cautiously because neither binding within the S9 incubation nor binding in blood was incorporated into the model. This limitation is particularly important for a lipophilic macrocyclic compound such as *ent*-verticilide, for which extensive nonspecific and protein binding could substantially affect the unbound concentration available for metabolism. Moreover, the intrinsic-clearance values were scaled using the human S9 scaling factor, rather than experimentally measured species-specific S9 scaling factors. The resulting hepatic-clearance estimates should therefore be considered comparative and hypothesis-generating rather than definitive quantitative predictions of in vivo clearance.

The predicted rat hepatic clearance of approximately 55 mL/min/kg exceeded the observed systemic clearance of 29.6 mL/min/kg following intravenous administration. This difference is not unexpected given the simplifying assumptions of the in vitro scaling model. In particular, omission of protein and incubation binding would be expected to overestimate clearance when only a small fraction of the compound is available for metabolism. Because systemic clearance includes the integrated effects of hepatic and extrahepatic elimination, the comparison does not independently establish the magnitude of hepatic clearance in vivo. Nevertheless, the observed systemic clearance, corresponding to approximately 42% of the rat hepatic blood flow of 70 mL/min/kg, supports appreciable in vivo elimination but was lower than the blood-flow-limited hepatic clearance predicted by the simplified well-stirred model, which did not incorporate plasma or S9 incubation binding.

Reaction phenotyping provided further evidence that P450-mediated metabolism contributes to the depletion of *ent*-verticilide in human hepatic preparations. Ketoconazole, a CYP3A4/5-selective inhibitor produced substantially greater preservation of *ent*-verticilide than inhibitors targeting CYP1A2, CYP2B6, CYP2C8, CYP2C9, CYP2C19, or CYP2D6. The response of the verapamil positive control supported effective CYP3A inhibition under the assay conditions. These findings identify CYP3A4/5 as the predominant P450 enzymes involved in *ent*-verticilide metabolism in pooled human liver microsomes. This finding is not surprising considering *ent*-verticilide is a large cyclic depsipeptide, as CYP3A4/5 has a large active site capable of accommodating such a substrate.^11^ However, ketoconazole inhibits both CYP3A4 and CYP3A5, and the relative abundance and activity of these enzymes in the pooled microsomal preparation were not established. The present experiment therefore does not distinguish the contributions of CYP3A4 and CYP3A5. Confirmation using recombinant CYP3A4 and CYP3A5, a CYP3A5-selective inhibitor, or P450-selective correlation analysis would strengthen the enzyme assignment. Furthermore, despite ketoconazole representing the only inhibitor observed to exhibit substantial inhibition of *ent*-verticilide metabolism, an appreciable amount of *ent*-verticilide depletion still occurred in the presence of ketoconazole (∼40% depletion). It is possible this observation may be due to a relatively long incubation period (60 minutes), given the extensive depletion of *ent*-verticilide in the control experiment (> 95% depletion). In addition, microsomes were preincubated with *ent*-verticilide prior to the addition of inhibitor, which may have reduced ketoconazole access to the enzyme. Finally, the possible contribution of other microsomal enzyme(s) that were not evaluated in the phenotyping study cannot be ruled out.

The metabolite-profiling results were consistent with the metabolic-stability and reaction-phenotyping findings. Eight putative metabolites were detected in human and rat liver S9 incubations containing NADPH and UDPGA. Their mass shifts were consistent predominantly with oxidative biotransformations, including mono- and dihydroxylation, combined hydroxylation and oxidation, oxidation, and desaturation. The absence of substantive formation of these signals in the buffer control containing cofactors but no liver S9 supports an enzymatic requirement for their formation. However, this control does not demonstrate cofactor dependence because it contained both NADPH and UDPGA.

M6, a putative monohydroxylated metabolite, was the most abundant metabolite signal in both human and rat liver S9 incubations. M2 was also prominent, particularly in rat S9, and the overall metabolite profiles were qualitatively similar between species. These findings suggest that human and rat hepatic preparations share several metabolic pathways for *ent*-verticilide. Nevertheless, differences in relative LC–MS response among the parent compound, C6, and M1–M8 preclude direct interpretation of relative peak areas as molar abundance. Consequently, the reported percentages should be considered semiquantitative signal distributions rather than quantitative estimates of the fraction of dose converted to each metabolite.

Although the reaction-phenotyping and metabolite-profiling results collectively support CYP3A-mediated oxidative metabolism, the present experiments cannot associate an individual P450 enzyme with a specific metabolite.

Reaction phenotyping measured only parent-compound depletion, whereas metabolite profiling was performed separately in liver S9 fractions without P450-selective inhibitors or recombinant enzymes. Therefore, CYP3A may contribute to the formation of one or several of M1–M8, but direct assignments cannot be made from the available data. Metabolite formation studies using P450-selective inhibitors, recombinant CYP3A4 and CYP3A5, or P450-enzyme panels would be required to establish specific P450–metabolite relationships.

No glucuronide conjugate was identified among M1–M8 despite the inclusion of UDPGA and alamethicin. This result suggests that glucuronidation did not generate a prominent detectable metabolite under the evaluated conditions, but it does not exclude a contribution of glucuronidation to *ent*-verticilide disposition. A glucuronide could have been formed below the reporting threshold, undergone in-source fragmentation, displayed poor ionization in positive-ion mode, or been unstable during sample processing or analysis. Evaluation in negative-ion mode, targeted searches for predicted glucuronide masses, and appropriate UGT-positive controls would provide a more definitive assessment. Lastly, because authentic metabolite reference standards were not available, metabolite structures and sites of biotransformation were considered tentative. The data support general biotransformation assignments, including hydroxylation, oxidation, and desaturation, but do not establish definitive sites of metabolism, regiochemistry, or stereochemistry.

Following intravenous administration in rats, *ent*-verticilide exhibited a steady-state volume of distribution of 4.86 L/kg, indicating distribution beyond the plasma compartment. Its systemic clearance of 29.6 mL/min/kg was below rat hepatic blood flow, and the terminal half-life was approximately 6.85 hours. The combination of a comparatively large distribution volume and non-flow-limited systemic clearance likely contributed to the persistence of measurable concentrations during the terminal phase.

After oral administration, *ent*-verticilide reached its observed maximum concentration at 1 hour, indicating that systemic absorption occurred relatively early after dosing. However, its absolute oral bioavailability was only approximately 10.5%. Low oral bioavailability may reflect incomplete gastrointestinal absorption, intestinal or hepatic first-pass metabolism, or a combination of these processes. Findings from our previous study, in which the P450 inhibitor 1-aminobenzotriazole (ABT) markedly increased the oral bioavailability of *ent*-verticilide, support a role for first-pass metabolism.^12^ Incorporation of reaction phenotyping in this study further confirms the contribution of CYP-mediated disposition of *ent*-verticilide. Additionally, the similar terminal half-lives following intravenous and oral administration provide no strong evidence of pronounced absorption-rate-limited elimination, although interpretation is constrained by the composite sampling design and the limited number of terminal sampling times.

Several limitations should be considered. Enzyme kinetic parameters were not obtained due to the lack of an authentic metabolite standard. Binding within the S9 and microsomal incubations was not measured, and the exploratory clearance predictions did not incorporate unbound fractions. The inhibitor-based reaction-phenotyping study did not independently distinguish CYP3A4 from CYP3A5 or establish the enzymes responsible for forming individual metabolites. Metabolite abundance was estimated from uncorrected LC–MS peak areas without authentic metabolite standards, and the proposed structures were not confirmed by synthesis, NMR, or comparison with reference standards. The presence of the C6 isomeric contaminant further complicated interpretation because it contributed materially to the total LC–MS signal and could potentially undergo metabolism independently of *ent*-verticilide.

Despite these limitations, the studies establish an initial drug metabolism and pharmacokinetics framework for *ent*-verticilide. The most important next steps are to determine unbound intrinsic clearance, distinguish CYP3A4 from CYP3A5, identify the P450 enzymes responsible for individual metabolite pathways, and confirm the structures of the major circulating or excreted metabolites. These studies will be important for predicting human disposition, evaluating susceptibility to CYP3A-mediated drug interactions, selecting appropriate nonclinical species, and guiding formulation or structural optimization of *ent*-verticilide.

## CONCLUSIONS

In conclusion, this study provides an initial characterization of the metabolism and pharmacokinetics of *ent*-verticilide. The findings demonstrate rapid hepatic metabolism, with CYP3A contributing prominently to oxidative biotransformation, and show that *ent*-verticilide undergoes multiple metabolic pathways across species. In rats, the compound exhibited extensive distribution and low oral bioavailability. Overall, these results establish a foundation for understanding the disposition of *ent*-verticilide and for guiding future studies supporting its continued development.

### Financial Support

This research was supported in part by the National Institutes of Health National Heart, Lung, and Blood Institute Grants [R35 HL144980] & [R01 HL183508] (B.C.K.), [R01 HL151223] (J.N.J., B.C.K., and W.S.A.), [R01 HL181098] (J.N.J., B.C.K., and W.S.A.), and [F31 HL164064] (M.P.T.), the American Heart Association Arrhythmia and Sudden Death Strategically Focused Research Network [Grant 19SFRN34830019] (B.C.K), and an American Heart Association Predoctoral Fellowship grant [24PRE1243081] (T.Q.D.), and a New Investigator Award from the American Association of Colleges of Pharmacy (R.D.C.).

### Data Availability

The authors declare that all the data supporting the findings of this study are available within the paper and its Supplemental Material.

### CRediT Authorship Contribution Statement

**Tri. Q Do**: Conceptualization, Data curation, Formal analysis, Funding acquisition, Investigation, Methodology, Project administration, Validation, Visualization, Writing — original draft, Writing — review and editing

**Rachel D. Crouch**: Conceptualization, Data curation, Formal analysis, Investigation, Methodology, Project administration, Validation, Visualization, Writing — original draft, Writing — review and editing

**Madelaine P. Thorpe**: Data curation, Formal analysis, Funding acquisition, Investigation, Methodology, Resources, Validation, Visualization, Writing — review and editing.

**Kevin M. Johnson**: Data curation, Formal analysis, Investigation, Methodology, Validation, Writing — original draft, Writing — review and editing

**Jeanne Rumsey**: Data curation, Formal analysis, Investigation, Methodology, Validation, Writing — original draft, Writing — review and editing

**Jeffrey N. Johnston**: Conceptualization, Data curation, Formal analysis, Funding acquisition, Investigation, Methodology, Project administration, Resources, Supervision, Validation, Visualization, Writing — review and editing

**Björn C. Knollmann**: Conceptualization, Data curation, Formal analysis, Funding acquisition, Investigation, Methodology, Project administration, Resources, Supervision, Validation, Visualization, Writing — review and editing

**Wendell S. Akers**: Conceptualization, Data curation, Formal analysis, Funding acquisition, Investigation, Methodology, Project administration, Resources, Supervision, Validation, Visualization, Writing — review and editing

## Supplemental Figures

**Figure S1:**
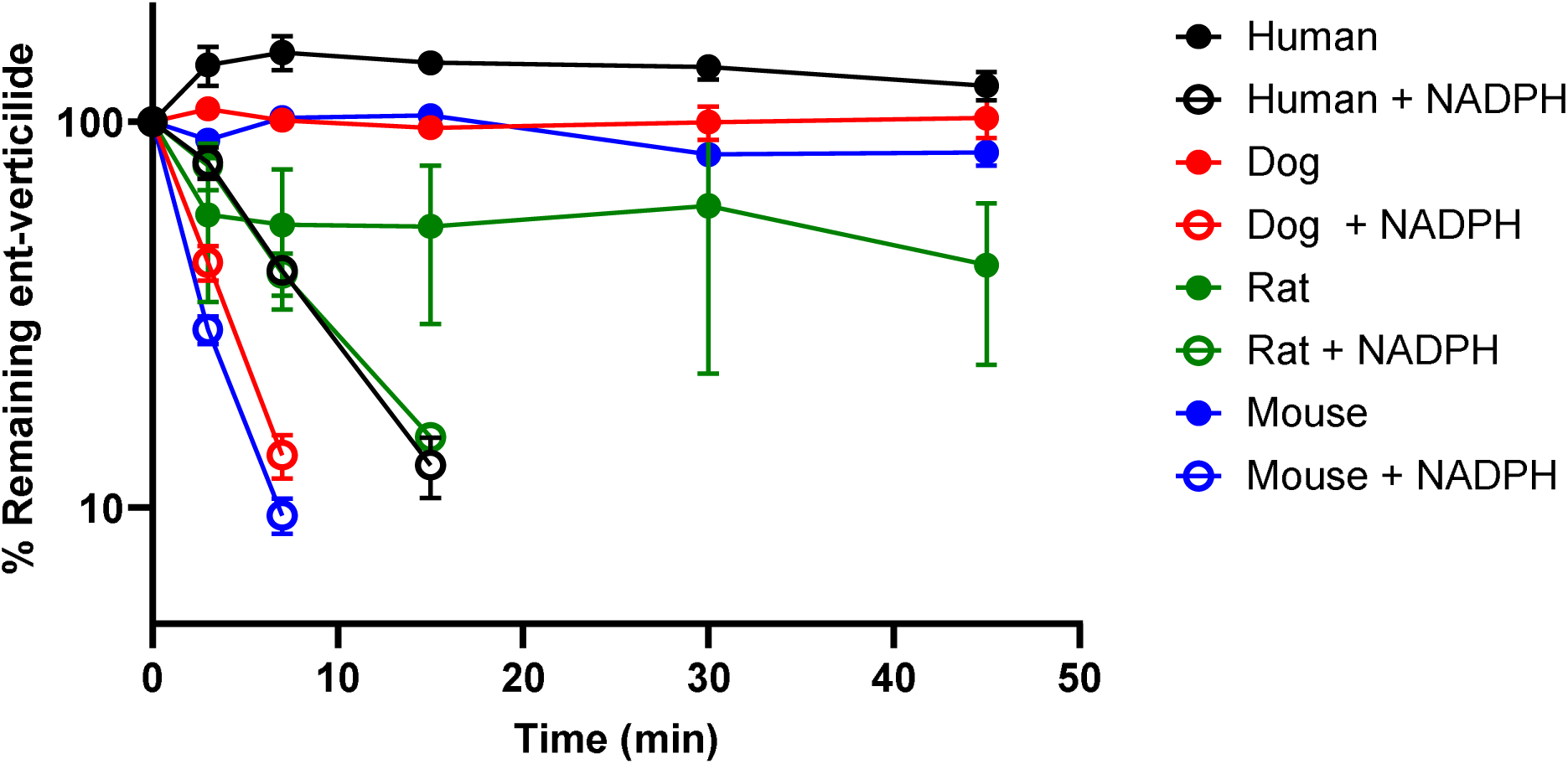
*ent*-Verticilide is Susceptible CYP-Mediated Metabolism in Mouse, Rat, Dog, and Human S9. Verapamil, a CYP3A4 substrate, was included as a positive control. Data = mean ± S.D. from a single experiment performed in triplicate. Data on the y-axis were natural-log transformed (ln) and were fitted using simple linear regression. **NADPH**: nicotinamide adenine dinucleotide phosphate.

**Figure S2.**
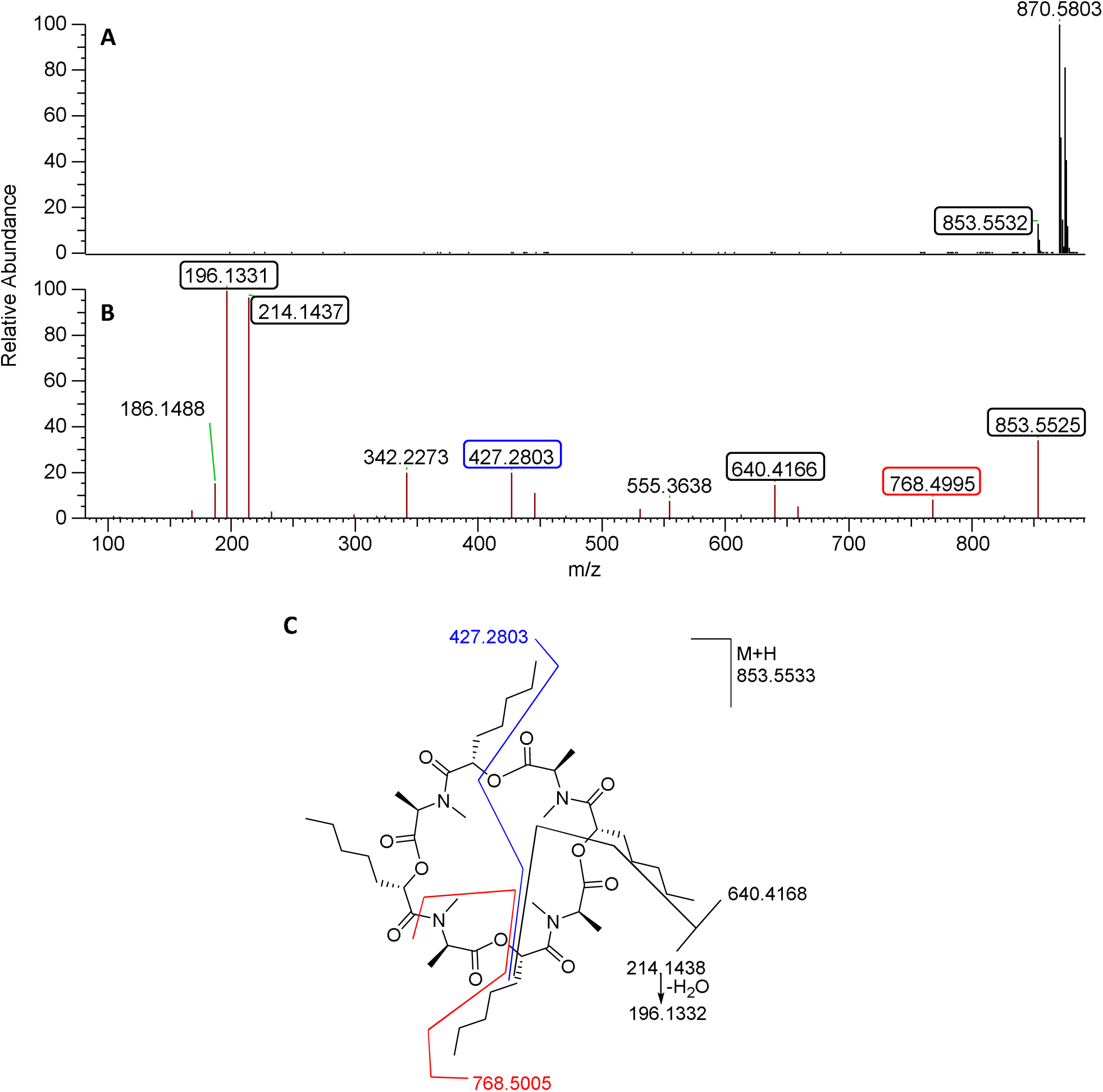
Full Scan MS and MS^2^ Spectra of ent-Verticilide (Positive ESI)

**Figure S3.**
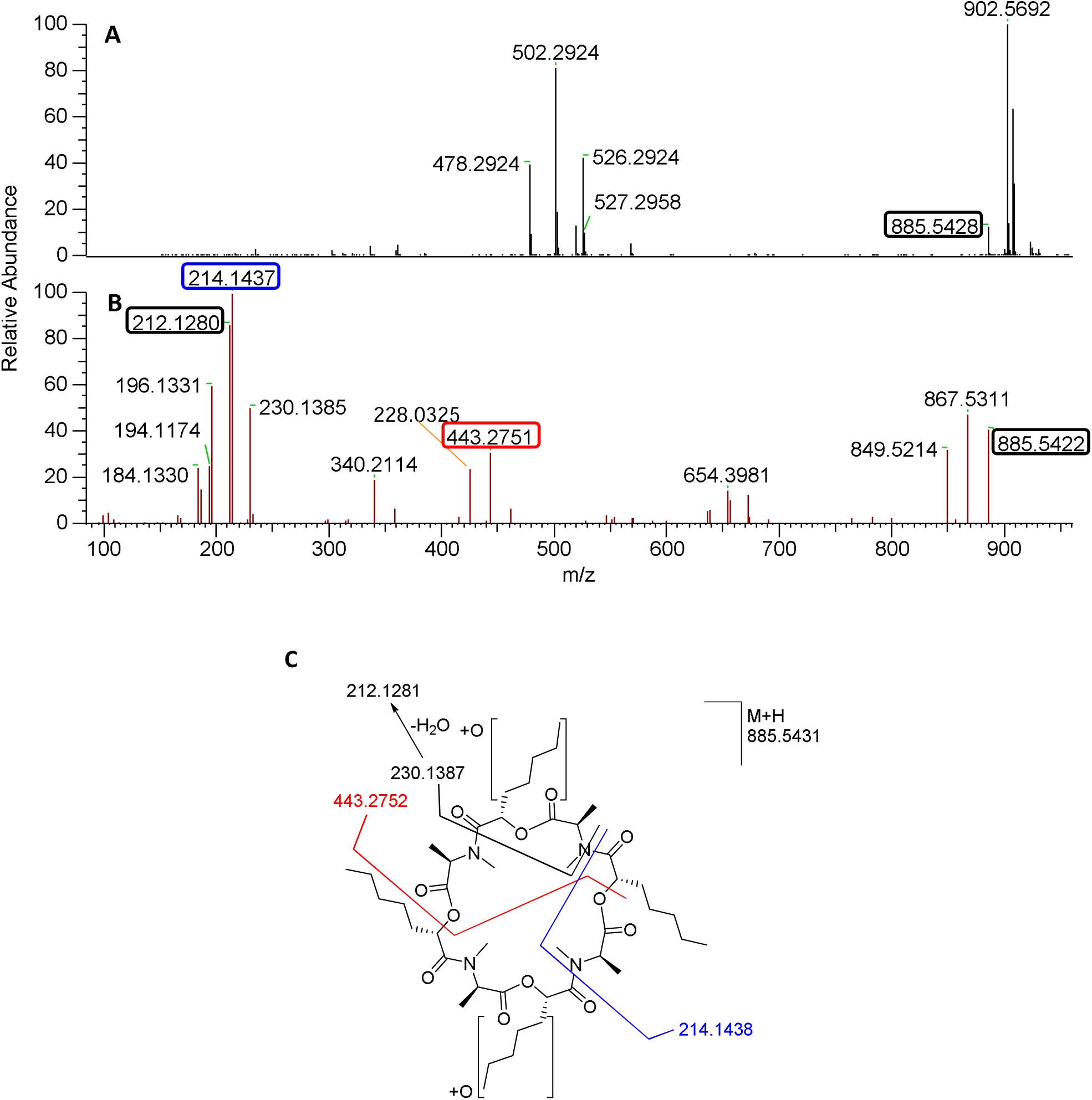
Full Scan MS and MS^2^ Spectra of M1 (Positive ESI).

**Figure S4.**
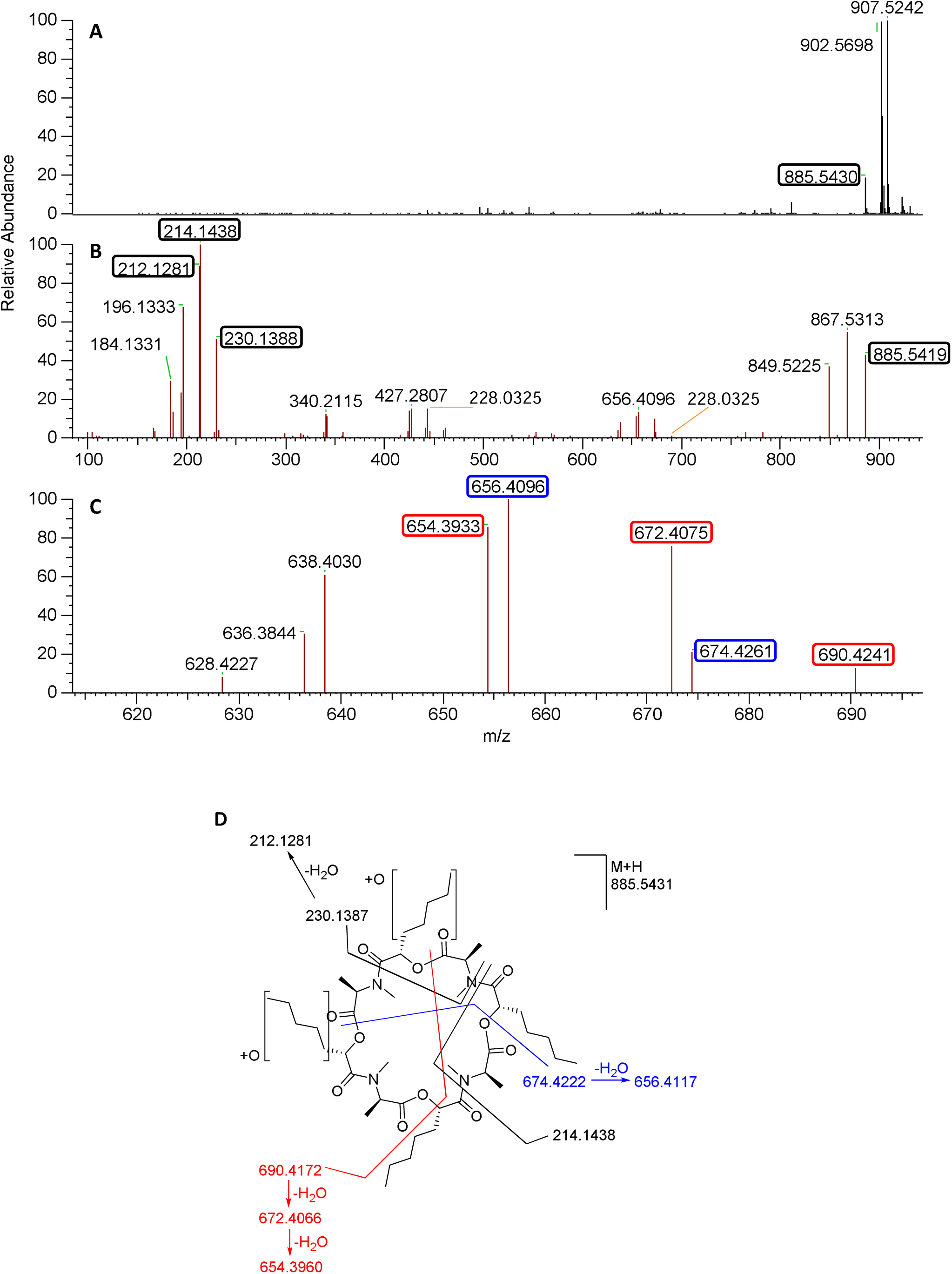
Full Scan MS and MS^2^ Spectra of M2 (Positive ESI).

**Figure S5.**
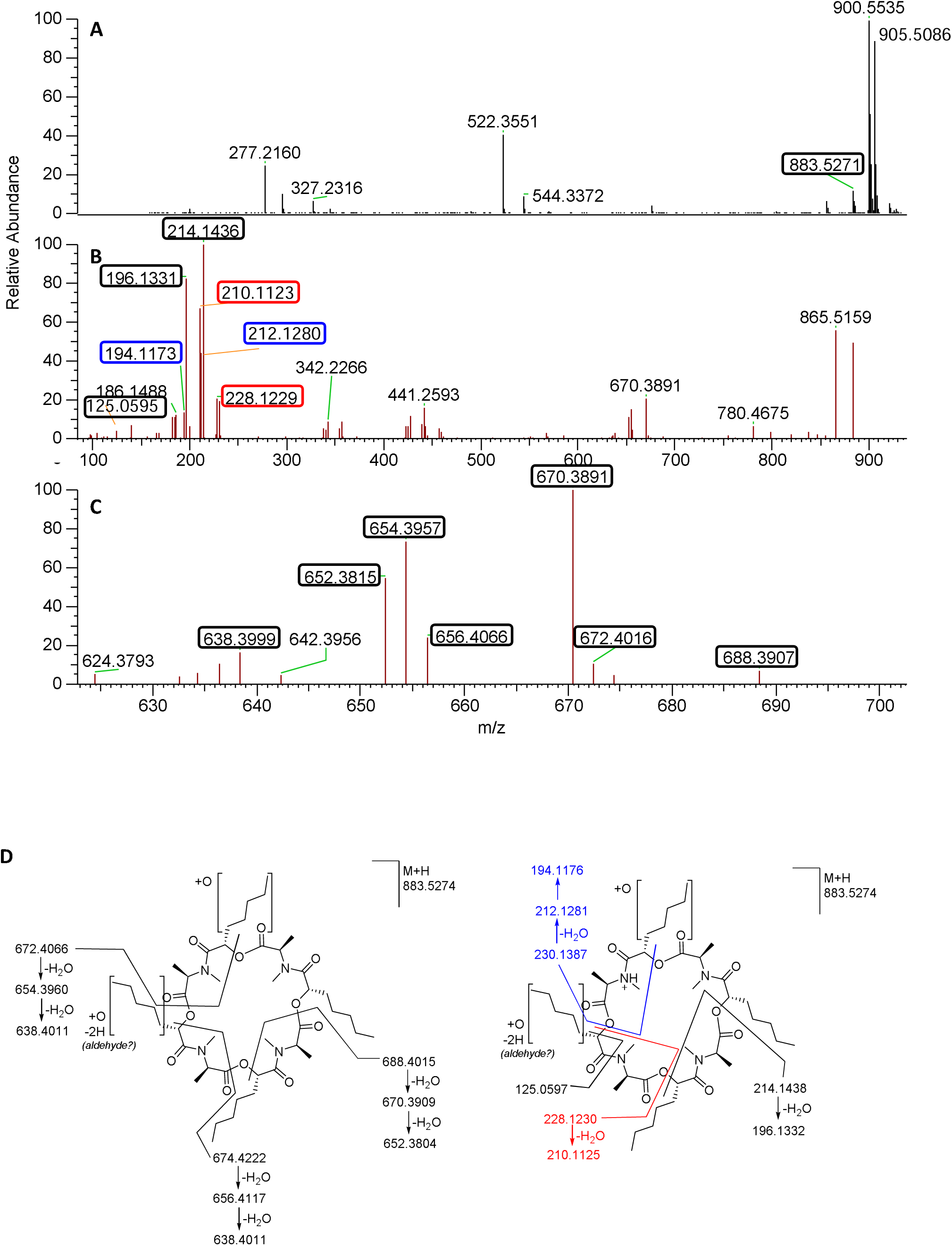
Full Scan MS and MS^2^ Spectra of M3 (Positive ESI).

**Figure S6.**
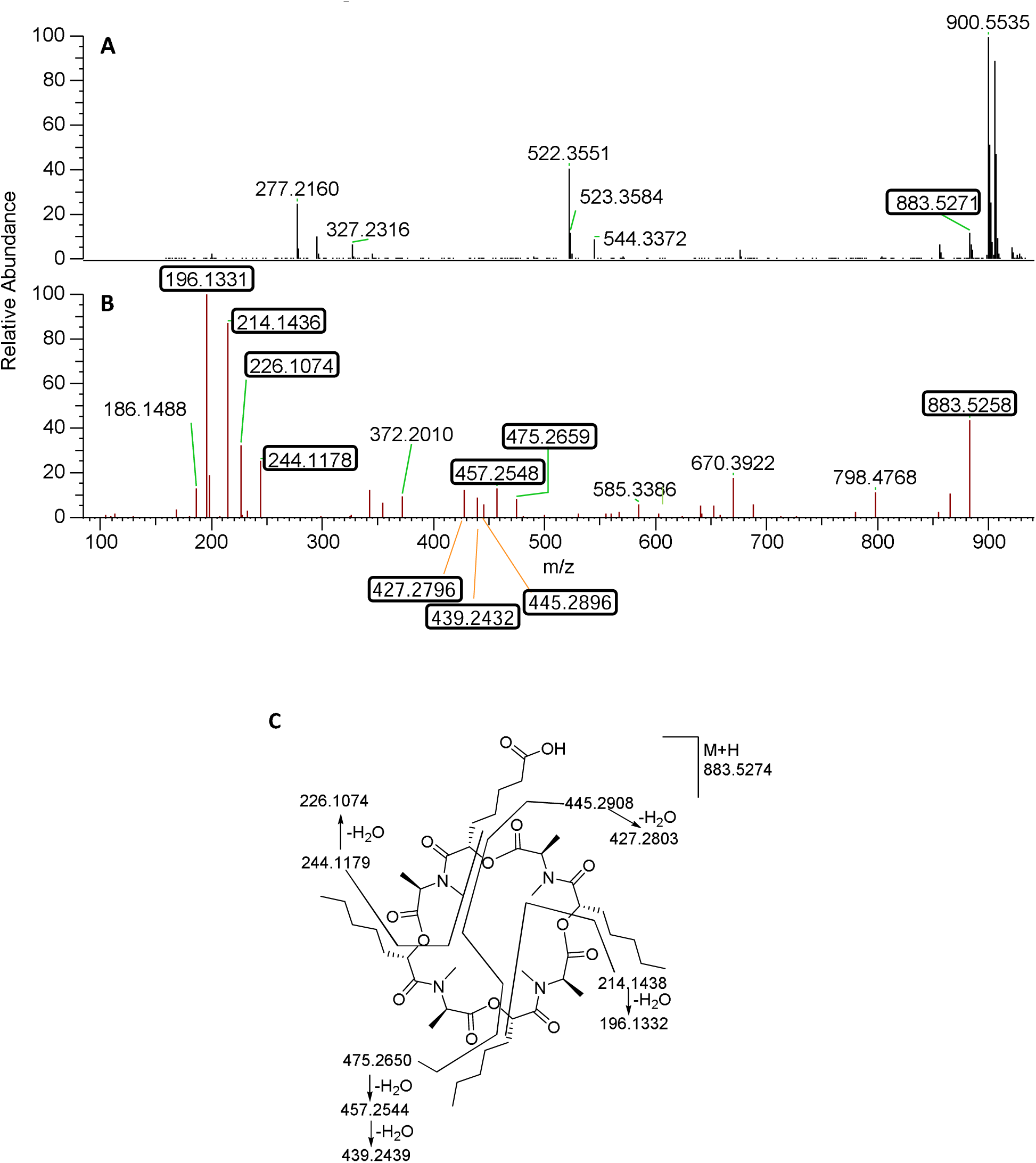
Full Scan MS and MS^2^ Spectra of M4 (Positive ESI).

**Figure S7.**
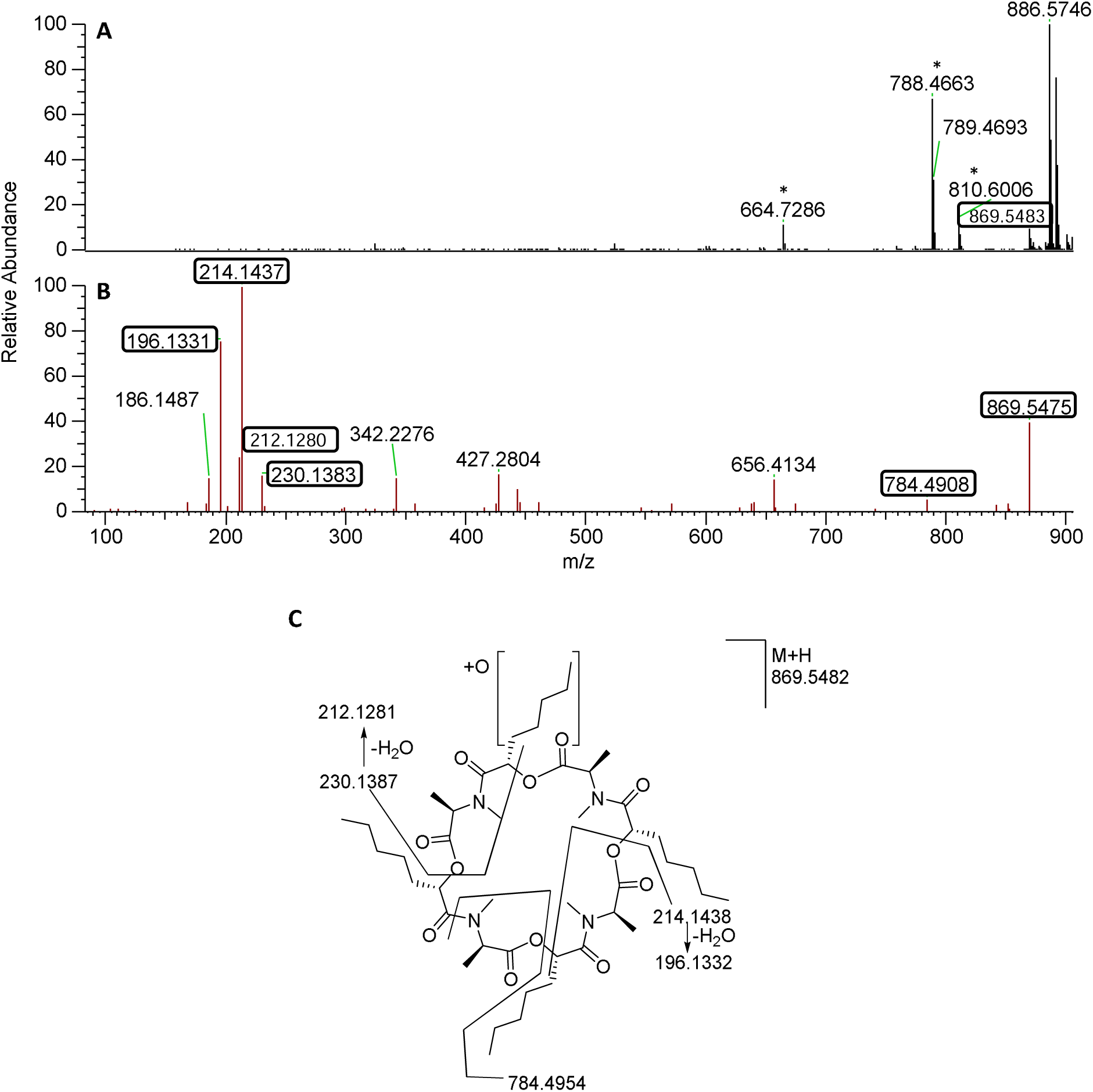
Full Scan MS and MS^2^ Spectra of M5 (Positive ESI).

**Figure S8.**
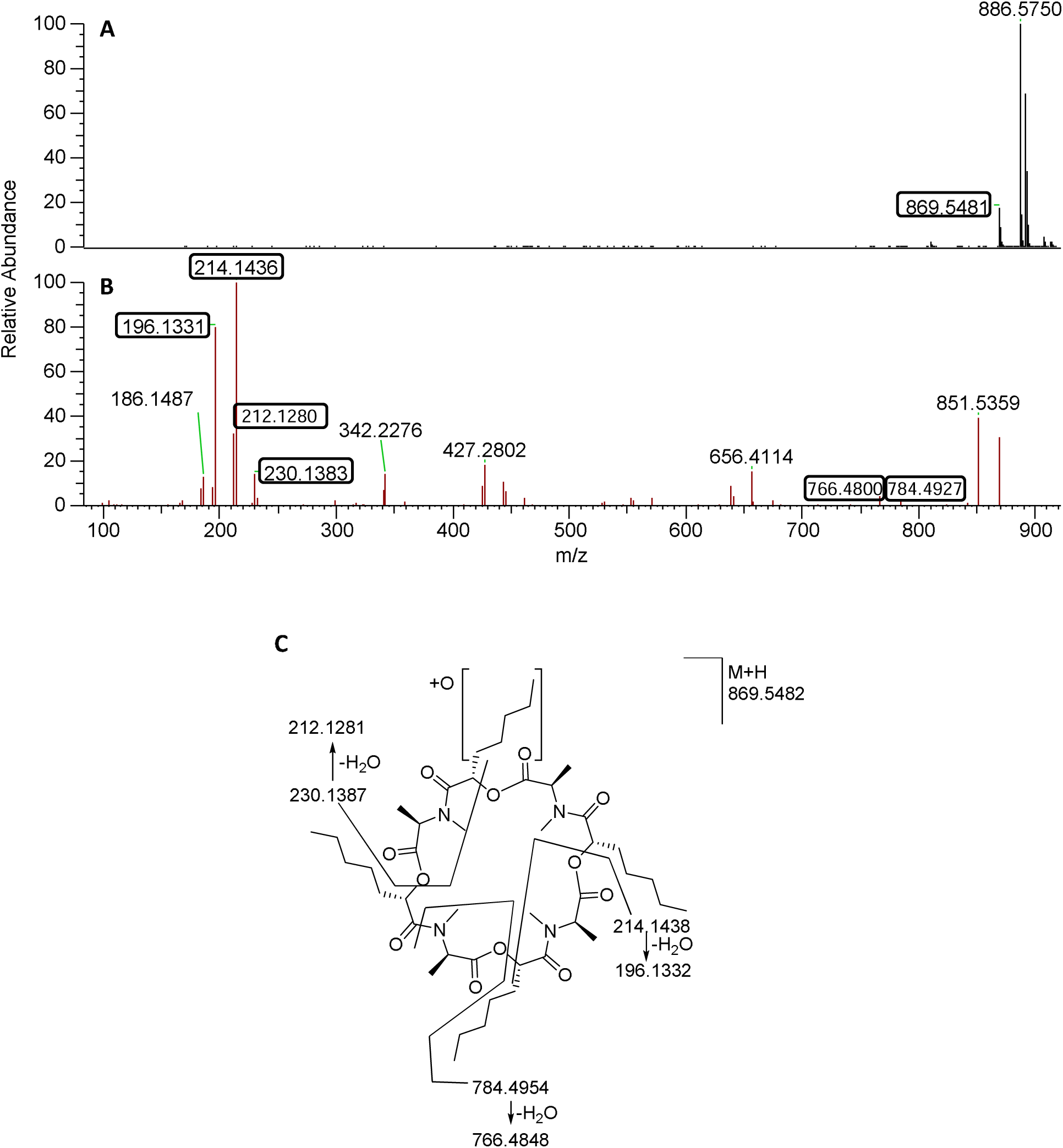
Full Scan MS and MS^2^ Spectra of M6 (Positive ESI).

**Figure S9.**
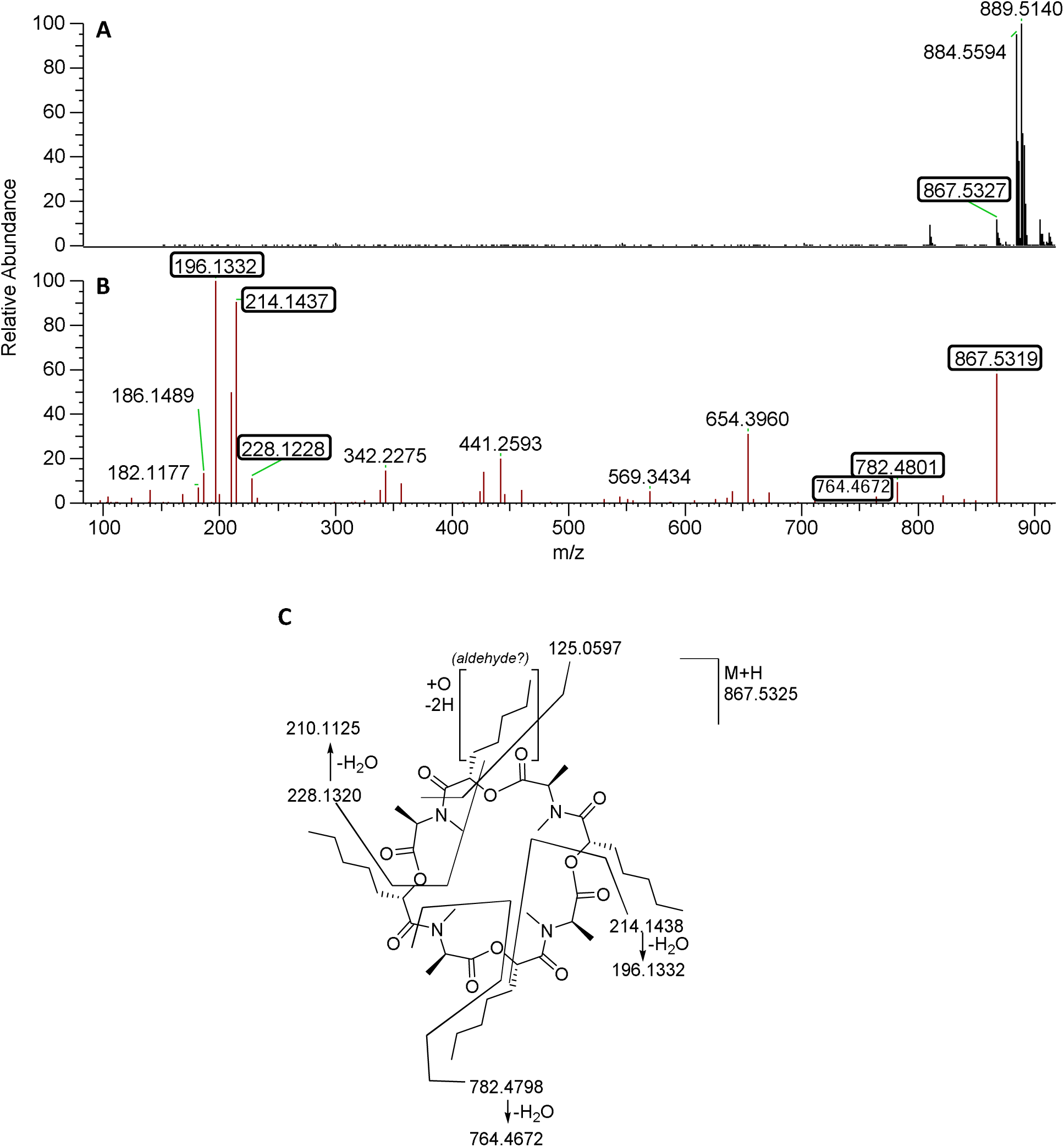
Full Scan MS and MS^2^ Spectra of M7 (Positive ESI).

**Figure S10.**
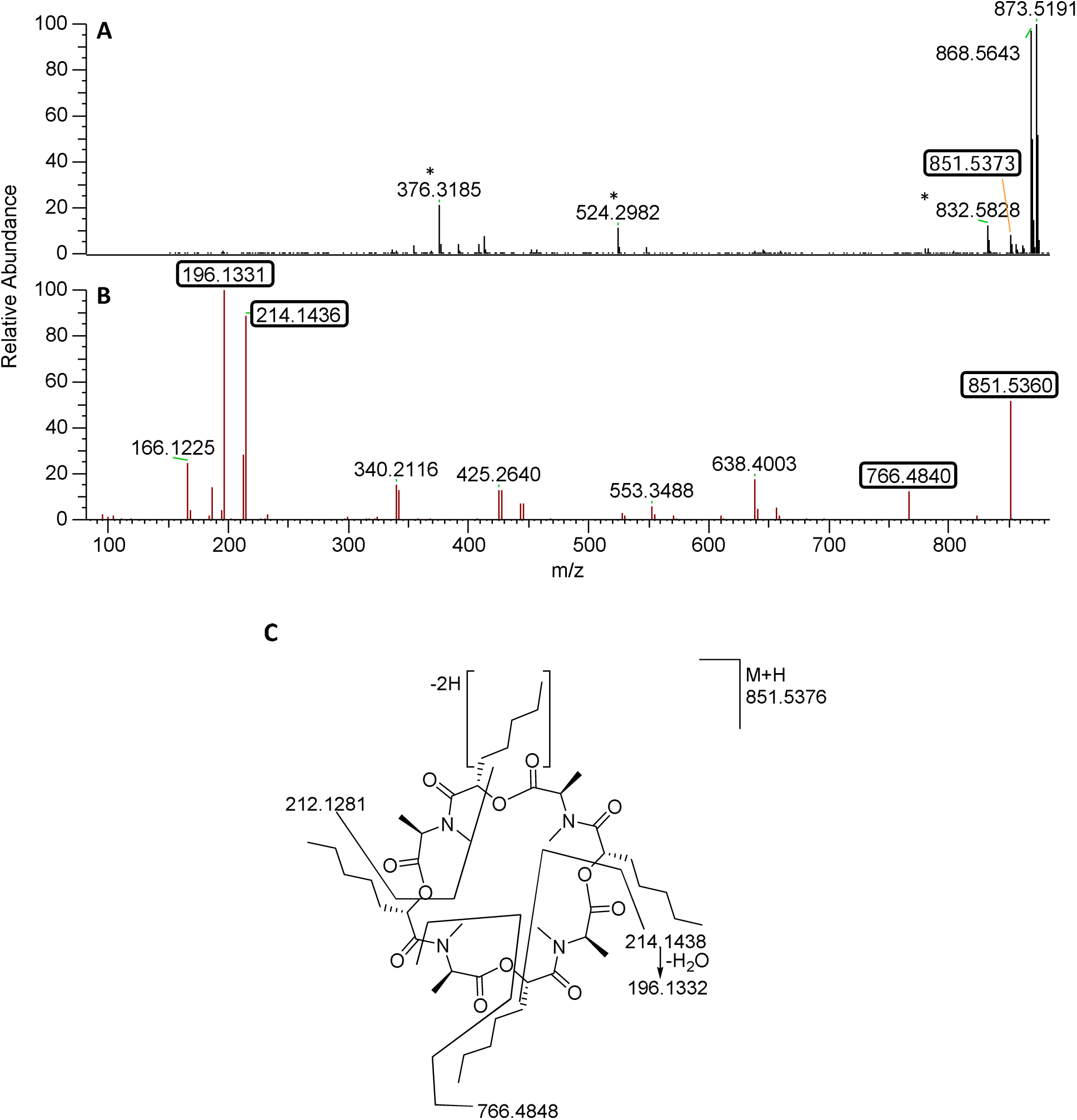
Full Scan MS and MS^2^ Spectra of M8 (Positive ESI).

## Supplemental Tables

**Table S1:** Inhibitors and the Corresponding Targeting Enzyme.

| **Chemical Inhibitors** | **Targeting Enzyme** | **Final Concentration (µM)** |
|---|---|---|
| α-naphthoflavone<sup>1</sup> | CYP1A2 | 1 |
| 2-phenyl-2-(1-piperidinyl)propane (PPP)<sup>2</sup> | CYP2B6 | 20 |
| Montelukast<sup>3</sup> | CYP2C8 | 2 |
| Sulfaphenazole<sup>4</sup> | CYP2C9 | 10 |
| (S)-(+)-N-3-benzylnirvanol<sup>5</sup> | CYP2C19 | 2 |
| Quinidine<sup>6</sup> | CYP2D6 | 1 |
| Ketoconazole<sup>7</sup> | CYP3A4/5 | 1 |

**Table S2.** HPLC-MS/MS Parameters for Analysis of *ent*-Verticilde, Verapamil, and Carbamazepine (Internal Standard).

| MS Parameters | *ent*-verticilide | verapamil | carbamazepine |
|---|---|---|---|
| Declustering potential (V) | 130 | 150 | 105 |
| Entrance potential (V) | 8 | 4 | 10 |
| Collision energy (V) | 39 | 36 | 25 |
| Collision cell exit potential (V) | 18 | 15 | 12 |
| Dwell time (ms) | 100 | 50 | 25 |
| Curtain gas (psi) | 35 | 35 | 35 |
| Collision-activated dissociation gas (psi) | 10 | 10 | 10 |
| Ion source gas 1 (psi) | 50 | 50 | 50 |
| Ion source gas 2 (psi) | 50 | 50 | 50 |

**Table S3:** Accurate Masses and Proposed Biotransformations of *ent*-Verticilide Metabolites.

| **Analyte** | **Proposed Biotransformation** | **Retention Time (min)** | **Observed [M+H]<sup>+</sup>** | **Theoretical [M+H]<sup>+</sup>** | **Mass Difference (ppm)** |
|---|---|---|---|---|---|
| M1 | Dihydroxylation (+32) | 9.0 | 885.5435 | 885.5431 | −0.5 |
| M2 | Dihydroxylation (+32) | 9.3 | 885.5430 | 885.5431 | 0.1 |
| M3 | Hydroxylation and oxidation (+30) | 9.5 | 883.5277 | 883.5274 | −0.3 |
| M4 | Hydroxylation and oxidation (+30) | 10.2 | 883.5273 | 883.5274 | 0.1 |
| M5 | Hydroxylation (+16) | 10.4 | 869.5482 | 869.5482 | 0.0 |
| M6 | Hydroxylation (+16) | 10.5 | 869.5482 | 869.5482 | 0.0 |
| M7 | Oxidation (+14) | 10.8 | 867.5320 | 867.5325 | 0.6 |
| M8 | Desaturation (−2) | 11.8 | 851.5378 | 851.5376 | −0.3 |
| *ent*-Verticilide | Parent compound | 12.3 | 853.5533 | 853.5533 | 0.0 |
| C6 | Isomeric contaminant | 12.6 | 853.5533 | 853.5533 | 0.0 |

## List of Non-standard Abbreviations

AUC: Area under the concentration versus time curve
DMSO: dimethyl sulfoxide
LC/MS: Liquid chromatography/mass spectrometry
LC-MS/MS: Liquid chromatography tandem mass spectrometry
P450: Cytochrome P450
PPP: 2-phenyl-2-(1-piperidinyl)-propane
MgCl_2_: Magnesium Chloride
NADPH: nicotinamide adenine dinucleotide phosphate tetrasodium salt
UDPGA: Uridine 5′-diphosphoglucuronic acid

